# Shu complex SWS1-SWSAP1 is required for mouse meiotic recombination in concert with the BRCA2 C terminus

**DOI:** 10.1101/238196

**Authors:** Carla M. Abreu, Rohit Prakash, Peter J. Romanienko, Ignasi Roig, Scott Keeney, Maria Jasin

## Abstract

Homology recognition and DNA-strand invasion ensure faithful homolog pairing and segregation during the first meiotic division^1^. RAD51 and DMC1 recombinases catalyze these steps^2^, with BRCA2 promoting their assembly into nuclear foci^3^. The recently identified human SWS1-SWSAP1 complex, related to the Shu complex in yeast, promotes RAD51 focus formation in cell lines^4,5^. We show here that mouse SWS1-SWSAP1 is critical for meiotic homologous recombination (HR) by promoting the assembly of RAD51 and DMC1 on early recombination intermediates. Absence of the complex perturbs meiotic progression in males and females and both sexes are sterile, although a fraction of meiocytes form crossovers. Remarkably, loss of the DNA damage checkpoint kinase CHK2 rescues fertility specifically in females without rescuing crossover numbers. Unlike the Shu complex, the BRCA2 C terminus (known to be required for RAD51 stabilization^6,7^) is dispensible for RAD51 and DMC1 focus formation. However, concomitant loss of the BRCA2 C terminus aggravates the meiotic defects in Shu mutant spermatocytes. These results point to a complex interplay of factors that ensure recombinase function and hence meiotic progression in the mouse.

---

Regulating the assembly, stabilization, and disassembly of nucleoprotein filaments of RAD51 and its meiosis-specific paralog DMC1 is critical for productive meiotic HR^2^. Transgenic mouse studies have shown that BRCA2, a multi-domain protein with several interaction sites for both recombinases^7-9^ is necessary for RAD51 and DMC1 focus formation during meiosis, such that BRCA2 loss causes meiotic arrest and sterility^3^. Other proteins are also expected to play a role in this process, including the canonical RAD51 paralogs, but understanding their meiotic role has been hampered by the embryonic lethality of mutants^10^. More recently, “Shu” complexes have been identified in several organisms; these interact with RAD51 and RAD51 paralogs and modulate RAD51^11^. Shu complexes are comprised of a defining member with a conserved Zn-coordinating motif and one or more members with RAD51-like structural motifs^4,5,11-13^. In budding yeast, mutations in any of the four subunits of the Shu complex <u>s</u>uppress the slow growth and <u>h</u>ydroxy<u>u</u>rea sensitivity (“Shu”) of *sgs1* and *top3* mutants^14^, but also promote meiotic HR by stabilizing Rad51 nucleoprotein filaments and promoting inter-homolog bias^13, 15, 16^. Structurally diverse Shu complexes found in *Caenorhabditis elegans* and *Schizosaccharomyces pombe* also play critical roles in meiotic HR^4,17^, but the role of the mammalian complex in this process is unknown.

To investigate the role of the mouse Shu complex, we disrupted *Sws1* (formally *Zswim7*) or *Swsap1* in fertilized eggs using TALE nuclease pairs directed to exon 1 downstream of the translation start site (Fig. 1a,b). From the several mutations obtained (Supplementary Table 1a,b), three frame-shift alleles for *Sws1* and two for *Swsap1* were selected for further analysis (Fig. 1a,b). Because results are similar for all alleles, most experiments in the main text focus on one mutant for each gene, *Sws1*^Δ*1(A)*/Δ*1(A)*^ and *Swsap1*^Δ*131*/Δ*131*^, hereafter *Sws1^-/-^* and *Swsap1^-/-^*, unless indicated otherwise in the figure legends. Surprisingly, unlike other RAD51 paralog knockout mice^10^, *Sws1^-/-^* and *Swsap1^-/-^* homozygous animals are viable, as are *Sws1^-/-^ Swsap1^-/-^* double mutants (Supplementary Table 2a,b). RT-PCR analysis using testis cDNA derived from *Sws1^-/-^* and *Swsap1^-/-^* mice confirmed the respective frame-shift alleles (Supplementary Fig. 1a). Mutant mice show no obvious gross morphological defects and have normal body weights (Supplementary Fig. 1b, 2a). However, neither male nor female *Sws1^-/-^* and *Swsap1^-/-^* mutants are fertile (Supplementary Table 3a). Testis weights from adult single and double mutants are 3- to 4-fold smaller than in control animals, and ovary weights are reduced 3- to 8-fold (Fig. 1c,d; Supplementary Fig. 1b and 2a). Notably, testis weights from mutant juveniles obtained before meiotic arrest (7.5 days postpartum, dpp) has occurred are similar to controls (Fig. 1c).

**Figure 1:**
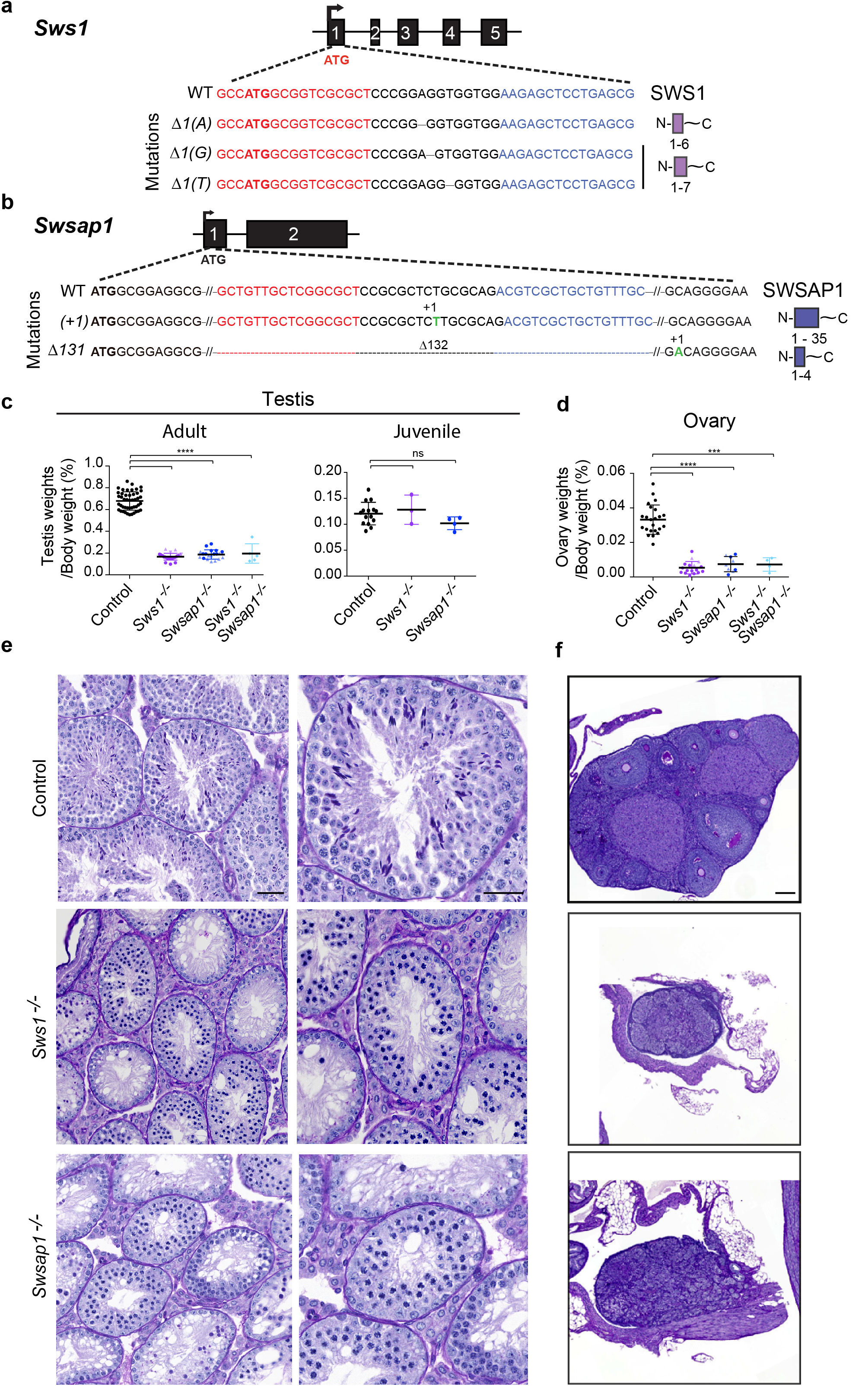
*Sws1* and *Swsap1* mutant mice show defective meiotic progression in both males and females. **(a,b)** *Sws1* (a) and *Swsap1* (b) genomic structure, TALEN target sequences (red and blue), mutations analyzed, and the predicted truncated proteins, if translated (shown on the right). **(c)** Testis to body weight ratios are reduced in adult single and double mutants, but not in juveniles. Adult mice: Control, n=45 (black circles); *Sws1^-/-^* alleles combined [Δ*1(A)*, n=20 (purple circles); Δ*1(G)*, n=4 (purple triangles); Δ*1(T)*, n=3 (purple inverted triangles)], *Swsap1^-/-^* alleles combined [Δ13l, n=10 (dark blue circles); *(+1)*, n=8 (dark blue triangles)], *Sws1^-/-^ Swsap1^-/-^* (Δ*1(A)*/(+1)), n=5 (light blue diamonds). Juvenile mice at 7.5 dpp: Control, n=15; *Sws1^-/-^*Δ*1(A)*, n=3; *Swsap1^-/-^*Δ*1A*, n=4. ns, not significant; ****, P<0.0001; Student’s t-test, two-tailed. **(d)** Ovary to body weight ratios are reduced in adult single and double mutants. Adult mice: Control, n=23; *Sws1^-/-^* [Δ*1(A)*, n=11; Δ*1(G)*, n=3; Δ*1(T)*, n=3], *Swsap1^-/-^* [Δ*131*, n=4; *(+1)*, n=5], *Sws1^-/-^Swsap1^-/-^* (Δ*1(A)* / (*+1*)), n=4. ***, P≤0.001; ****, P≤0.0001; Student’s t-test, twotailed. **(e)** Shu mutant spermatocytes arrest at pachynema. Sections were stained with periodic acid-Schiff (PAS). Scale bars, 100 μm and 50 μm for left and right columns, respectively. Control, *Sws1^-/-^, Swsap1^-/-^*, n≥3. **(f)** Ovaries from Shu mutant adults lack follicles. Sections were stained with PAS. Scale bar, 500 μm. Control, *Sws1^-/-^, Swsap1^-/-^*, n≥3.

In testis sections from adult Shu-mutant mice, seminiferous tubules have substantially reduced cellularity and are devoid of post-meiotic germ cells (Fig. 1e and Supplementary Fig. 1c). Spermatocytes appear to arrest during mid-pachynema, possibly at stage IV of the seminiferous epithelial cycle^18^. TdT-mediated dUTP nick end-labeling (TUNEL) demonstrates widespread apoptosis (Supplementary Fig. 2b). Apoptosis in mutant juvenile testes at 7.5 dpp is rarely observed, as in controls, suggesting that pre-meiotic cells are not affected (Fig. 1c and Supplementary Fig. 2c). Adult ovary sections from mutants lack follicles at any developmental stage (Fig. 1f and Supplementary Fig. 2d). At 3 dpp, ovaries stained for c-Kit, a marker for diplotene and dictyate stage oocytes in primordial and primary follicles, have significantly reduced oocyte numbers in *Sws1^-/-^* and *Swsap1^-/-^* mice, with some oocytes appearing to be apoptotic (Supplementary Fig. 2e). Together, our data demonstrate that SWS1 and SWSAP1 are essential for meiotic progression in both male and female mice.

Testes and ovaries from Shu-mutant mice resemble those of HR- and synapsis-defective mutants, such as *Dmc1^-/-^* and *Sycp1^-/-^* (Ref.^19-22^). To test if the Shu complex is required for HR and/or synapsis, we analyzed the synaptonemal complex (SC), a tripartite proteinaceous structure that forms between the homolog axes as they pair, by immunostaining surface-spread spermatocytes for the SC central region (SYCP1) and axial/lateral elements (SYCP3)^1^. Spermatocytes were also stained for the testis-specific histone H1 variant (H1t), which specifically labels cells at mid-pachynema and beyond^23^. H1t-positive spermatocytes are significantly reduced in mutant testes, indicating an early-pachytene arrest that is bypassed in only a fraction of cells (Fig. 2a,b and Supplementary Fig. 3a,b). The ability of some cells to progress contrasts with *Dmc1^-/-^*, in which H1t-positive cells are absent^24^. Unlike later stages, early meiotic prophase cells at leptonema and zygonema are increased in Shu single- and doublemutant mice. Synaptic abnormalities begin to be observed at early zygonema, such that chromosomes are seen with long axes but no synapsis, which becomes more pronounced by late zygonema, where chromosomes with fully formed axes are found with little or no synapsis (“early- and late zygonema-like”, respectively; Fig. 2b,c and Supplementary Fig. 3b,d). In contrast to wild-type pachytene cells in which all of the homologs are typically synapsed, the majority of mutant cells are abnormal at this stage, displaying unsynapsed or partially synapsed chromosomes and frequent synapsis between non-homologous chromosomes (“pachynemalike”; Fig. 2b,c and Supplementary Fig. 3b,d,e). While severe, the synaptic defects are not as profound as reported for *Dmc1^-/-^* mutants^19,20^. Synaptic defects are also observed in the sex chromosomes, with less than half of mutant spermatocytes at early pachynema having synapsed XY pairs (Supplementary Fig. 3c). Mutant cells with autosomal synapsis defects are more likely to also have unsynapsed XY pairs, whereas those with full autosomal synapsis typically have synapsed XY pairs. The few cells that reach mid-pachynema tend to have fewer synaptic abnormalities (Fig. 2b and Supplementary Fig. 3b,d).

**Figure 2:**
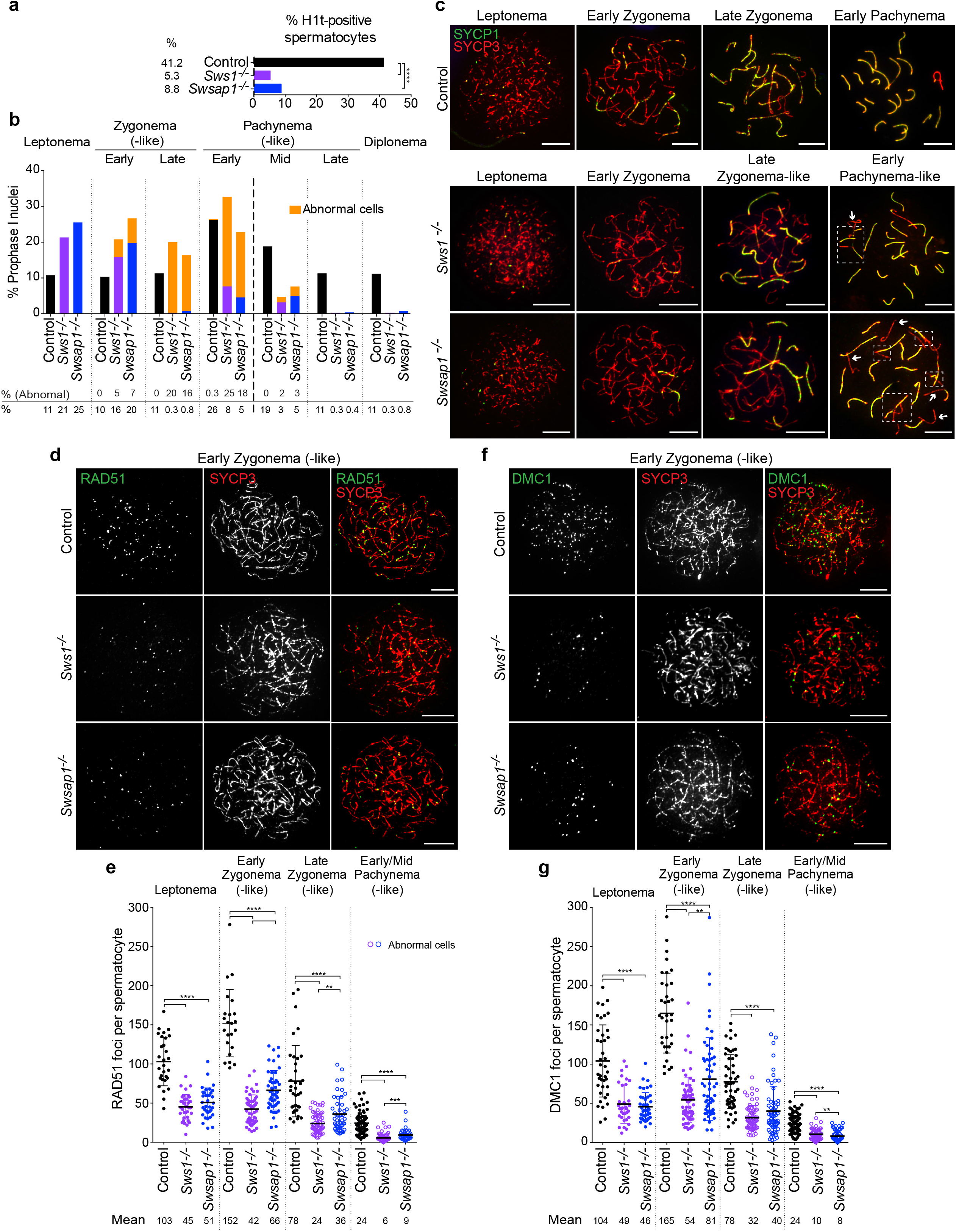
SWS1 and Swsap1 are required for normal homolog synapsis, DSB repair, and RAD51 and DMC1 focus assembly during meiotic HR. **(a)** Histone H1t staining indicates that few spermatocytes in *Sws1* and *Swsap1* mutants progress to mid-pachynema and beyond. Total number of mid-pachytene, late-pachytene, and diplotene spermatocytes divided by the total number of spermatocytes analyzed from adult testes. Mice: Control, n=6; *Sws1^-/-^, Swsap1^-/-^*, n=3. ****, P≤0.0001; Fisher’s exact test, two-tailed. **(b,c)** *Sws1* and *Swsap1* mutants show altered meiotic progression and abnormal chromosome synapsis. Percentage of spermatocytes in each of the indicated meiosis prophase I stages in **b** with representative images in **c**. *Sws1^-/-^* and *Swsap1^-/-^* cells at leptonema are indistinguisable from controls. At later stages abnormal cells with synaptic defects are observed; because synapsis is abnormal these stages are appended with the word “-like” (orange bars). At early zygonema, chromosomes begin to synapse in the majority of mutant cells, althought the number of synaptic stretches are usully reduced; however, delayed synapsis onset is apparent in a subset of mutant cells, as indicated by the lack of SYCP1 stretches. At late zygonema, abnormal cells with fully formed chromosome axes (SYCP3) but little or no synapsis (SYCP1) predominate. Fully-synapsed chromosomes with thicker and shorter SYCP3 axes characterize early pachynema, indicative of chromatin condensation, however, abnormal cells with unsynapsed chromosomes and chromosomes with non-homologous synapsis and/or partial asynapsis are frequent in the mutants. Mid-pachytene (H1t-positive) abnormal cells contain unsynapsed chromosomes, broken bivalents, or parts of chromosomes involved in non-homologous synapsis. Scale bars, 10 μm. Boxes in c in early pachytene-like cells highlight non-homologous synapsis and arrows indicate unsynapsed chromosomes. **(d-g)** RAD51 and DMC1 focus counts are reduced in *Sws1* and *Swsap1* mutant spermatocytes. Representative chromosome spreads from adult mice at early zygonema (-like) are shown in **d,f** with focus counts for all stages in **e,g**. n=3. Error bars, mean±s.d. Scale bars, 10 μm. Each circle in **e,g** indicates the total number of foci from a single nucleus. Solid circles in **e, g**, normal cells. Open circles in e,g, cells with abnormal synapsis. **, P≤0.01; ***, P≤0.001; ****, P≤0.0001; Mann-Whitney test, one-tailed.

Synapsis defects in Shu-mutant spermatocytes could reflect meiotic HR defects, as the human Shu complex promotes HR in cultured cells^5^. Indeed, *Sws1^-/-^* and *Swsap1^-/-^* spermatocytes display an ~2-fold reduction in RAD51 and DMC1 focus numbers at leptonema (Fig. 2d-g and Supplementary Fig. 3f,g). RAD51 and DMC1 focus numbers increase substantially by early zygonema in control cells, but remain low in mutant cells. At later stages, focus numbers progressively decrease in all genotypes, including the mutants. Shu double-mutant spermatocytes have similarly reduced RAD51 and DMC1 focus numbers at all stages (Supplementary Fig. 3f,g).

Sterility can occur in mutants where a similar reduction in RAD51 and DMC1 foci is attributable to fewer DSBs^25^. To rule out effects of Shu complex loss on DSB formation and/or their resection, we quantified chromatin-bound γH2AX^26^, a marker for DSBs, and foci of MEIOB, a meiosis-specific, single-stranded DNA (ssDNA)-binding protein^27,28^. At leptonema and early zygonema, γH2AX levels are indistinguishable from controls (Supplementary Fig. 4a,b), suggesting that DSB formation is unaffected. Further, there are more MEIOB foci at these stages in Shu mutant spermatocytes (2.4-fold at leptonema and 1.2-fold at early zygonema; Fig. 3a,b and Supplementary Fig. 4c), indicating an increase in the number of end-resected intermediates that are unable to stably assemble RAD51 and DMC1. Notably, the increase in MEIOB foci is not as great as in *Dmc1^-/-^* spermatocytes (3.3- and 1.8-fold, respectively). We interpret these findings to indicate that DSBs are formed in normal numbers and are resected, to be initially bound by MEIOB, in *Sws1^-/-^* and *Swsap1^-/-^* mutants, but that the mouse Shu complex fosters the stable assembly of RAD51 and DMC1 nucleoprotein filaments during meiotic HR, which in turn promotes homolog synapsis.

**Figure 3:**
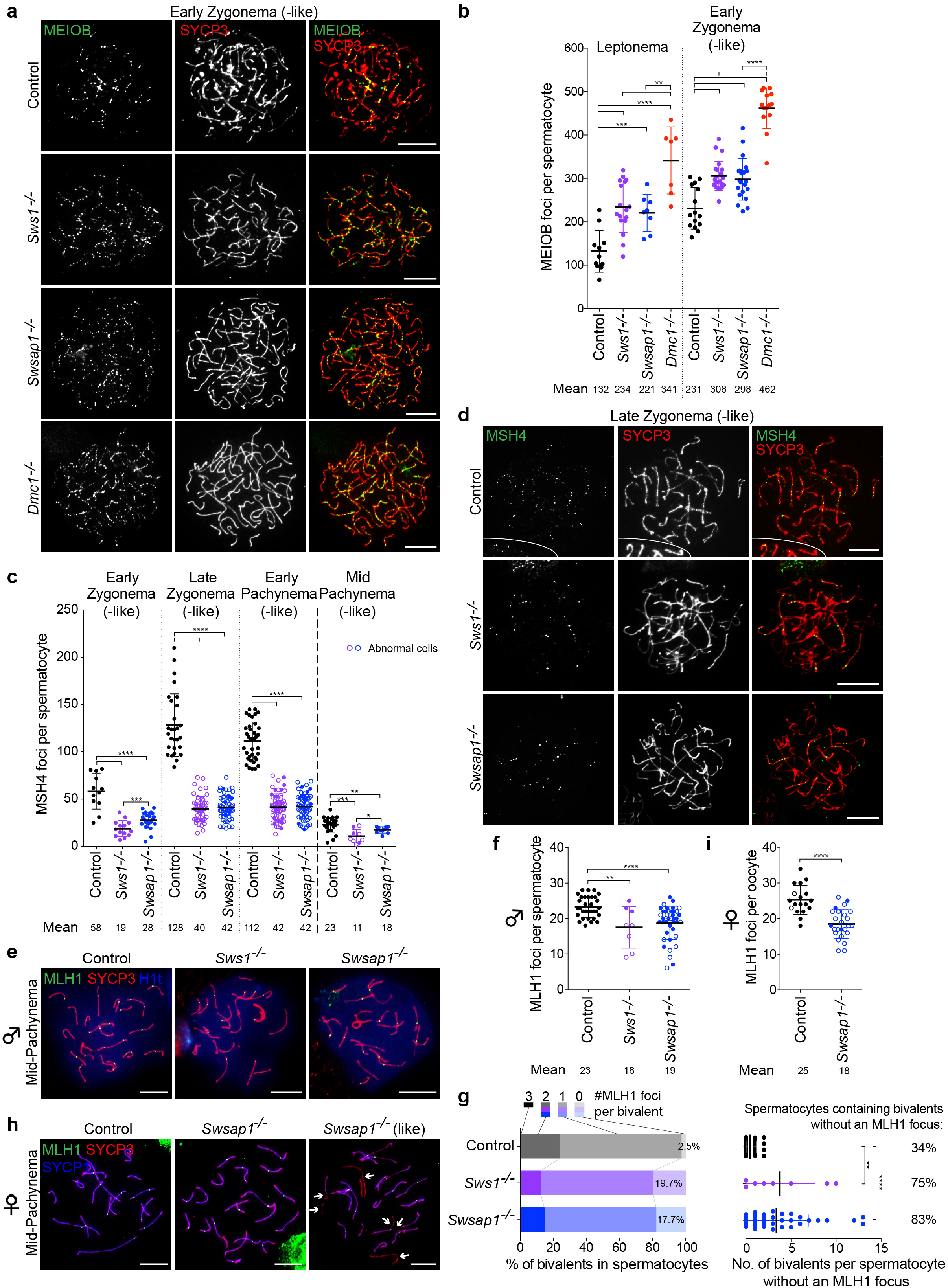
SWS1 or Swsap1 loss causes accumulation of resected DNA intermediates early in meiosis and defects at later meiotic stages. **(a,b)** MEIOB focus counts are increased in *Sws1^-/-^* and *Swsap1^-/-^* spermatocytes, but not as much as in *Dmc1^-/-^* spermatocytes. Representative chromosome spreads at early zygonema (-like) from adult mice in a with focus counts at indicated stages in **b**. Mice: Control, n=2; *Sws1^-/-^, Swsap1^-/-^*, n=3; *Dmc1^-/-^*, n=1. **(c,d)** MSH4 focus counts are reduced in *Sws1^-/-^* and *Swsap1^-/-^* spermatocytes. Focus counts at indicated stages in **c** with representative images of chromosome spreads at late zygonema (-like) in **d**. Mice: Control, n=2; *Sws1^-/-^, Swsap1^-/-^*, n=3. **(e,f)** MLH1 focus counts are reduced on average by ~20% in *Sws1^-/-^* and *Swsap1^-/-^* spermatocytes at mid-/late pachynema (-like). Representative chromosome spreads at mid-pachynema in e with focus counts in **f**. Mice: Control, *Swsap1^-/-^*, n=2; *Sws1^-/-^*, n=1. **(g)** Although most *Sws1^-/-^* and *Swsap1^-/-^* bivalents have at least one MLH1 focus (left), most spermatocytes have at least one bivalent which lacks a focus. Left, percentage of bivalents in mid-/late pachynema (-like) with indicated number of MLH1 foci per bivalent. Right, number of bivalents without an MLH1 focus per spermatocyte; total percentage of spermatocytes containing bivalents without an MLH1 focus is indicated. **(h,i)** MLH1 focus counts are reduced on average by ~30% in *Sws1^-/-^* and *Swsap1^-/-^* oocytes. Representative chromosome spreads at mid-pachynema (-like) from mice at embryonic day 18.5 in **h** with focus counts in **i**. n=1 for both genotypes. While a fraction of mutant oocytes show completely synapsed bivalents (solid circles in **i**), the majority have chromosomal synapsis defects (open circles), i.e., fully unsynapsed and/or partially synapsed bivalents. In controls, a small fraction of oocytes also have one or two chromosome ends that are not fully synapsed. Each circle in **b,c,f,i** indicates the total number of foci from a single nucleus. Solid circles, normal cells. Open circles, cells with abnormal synapsis. Error bars in **b,c,f,g,i**, mean±s.d. Scale bars in **a,d,e,h**, 10 μm. *, *P*≤0.05 in c; **, *P*≤0.01 in **b,c,f,g**; ***, P≤0.001 in **b,c**; ****, *P*≤0.0001 in **b,c,f,i,g**; Mann-Whitney test, one-tailed.

Given the early meiotic prophase I defects in Shu-mutant spermatocytes, we expected that HR is impaired later as well. Consistent with defects in DSB repair, mutant spermatocytes at early pachynema display γH2AX on autosomes, which is even more evident in early pachytene-like cells with synapsis defects (Supplementary Fig. 4d). γH2AX mostly disappears from autosomes in control cells and remains concentrated in the unsynapsed XY chromatin forming the sex body^26^. In contrast, autosomal γH2AX in Shu-mutant cells is accompanied by defects in sex body formation/maturation, especially in those cells with a high degree of autosome asynapsis (Supplementary Fig. 4d,e). Interestingly, the rare mid-pachytene Shu-mutant spermatocytes display less autosomal γH2AX than those at early pachynema and occasionally mature sex bodies (Supplementary Fig. 4f), suggesting that some mutant cells do not trigger the pachytene checkpoint due to greater proficiency in DSB repair. However, γH2AX remnants are still observed in these *Sws1^-/-^* and *Swsap1^-/-^* “escapers”, in agreement with evidence suggesting that the pachytene checkpoint tolerates some unrepaired DSBs^29^.

Because a small fraction of Shu-mutant spermatocytes are apparently repair-proficient and progress to mid-pachynema, we asked whether mutant cells could form later HR intermediates. MSH4 stabilizes DNA-strand exchange intermediates, some of which will become crossovers, whereas MLH1 specifically marks crossovers^1^. Mutant spermatocytes have 2- to 3fold fewer MSH4 foci from early zygonema to early pachynema, proportional to the earlier reduction in the RAD51 and DMC1 foci (Fig. 3c,d). Remarkably, however, MLH1 foci are reduced on average only ~20% in mid-pachytene cells, and the majority of bivalents have at least one MLH1 focus even though most cells have one or more chromosome pairs lacking a focus (Fig. 3e-g). Considering the number of MLH1 foci, it is striking that none of the mutant cells at early pachynema have MSH4 focus numbers within one standard deviation of the mean in control cells (Fig. 4c). These results suggest that crossover homeostasis^30,31^ operates in the Shu mutants.

**Figure 4:**
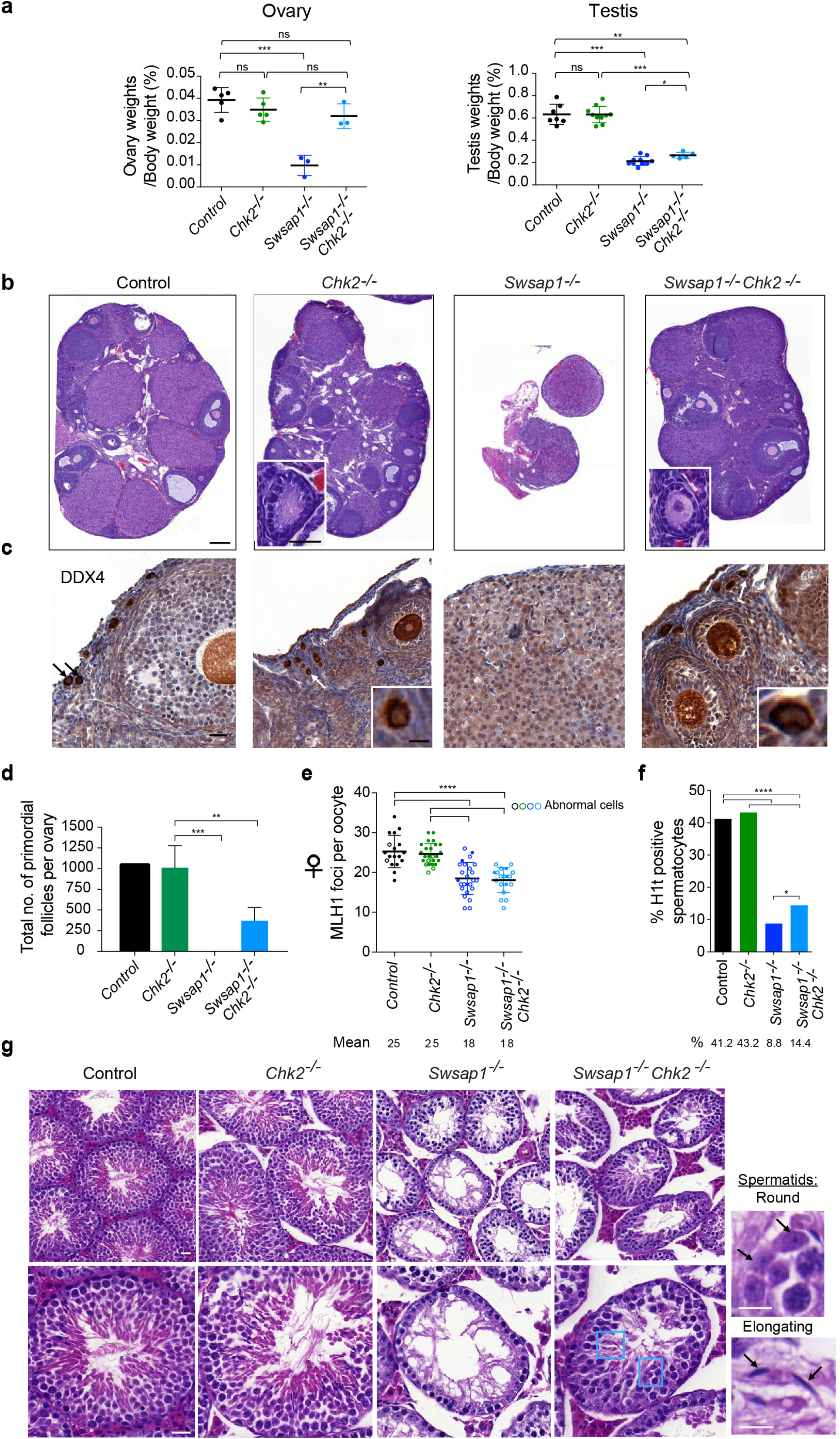
*Chk2* mutation rescues fertility of *Swsap1* mutant females, but not males, without rescuing synapsis defects or MLH1 focus numbers. **(a)** Ovary to body weight ratios of *Swsap1^-/-^ Chk2^-/-^* adult mice are similar to controls, but testis to body weight ratios remain significantly reduced. Female mice: Control, *Chk2^-/-^*, n=5; *Swsap1^-/-^, Swsap1^-/-^ Chk2^-/-^*, n=3. Male mice: Control, n=7; *Chk2^-/-^, Swsap1^-/-^*, n=10; *Swsap1^-/-^ Chk2^-/-^*, n=5. *, *P*≤0.05; **, *P*≤0.01; ***, *P*≤0.001; Student’s t-test; two-tailed. **(b-d)** Adult *Swsap1^-/-^ Chk2^-/-^* ovaries have follicles at various stages of oocyte development, unlike *Swsap1^-/-^* ovaries. Primordial follicles are highlighted in **b** in insets and by arrows in c and are quantified in d. Sections were stained with hematoxylin and eosin (H&E) in **b**, and with hematoxylin and an antibody for DDX4/Vasa in c. Scale bar, 500 μm and 50 μm insets. Mice: Control, n=1; *Chk2^-/-^, Swsap1^-/-^*, *Swsap1^-/-^ Chk2^-/-^*, n=3. *P* values in **d** were obtained using a oneway ANOVA test (Kruskal Wallis); **, *P*≤0.01; ***, *P*≤0.001. **(e)** MLH1 foci are reduced in *Swsap1^-/-^ Chk2^-/-^* oocytes from mice at embryonic day 18.5 similar to the level observed in *Swsap1^-/-^* oocytes, and synaptic defects are also observed in the majority of *Swsap1^-/-^ Chk2^-/-^* oocytes as also seen in *Swsap1^-/-^* mice (open circles). Mice: Control, *Swsap1^-/-^*, from Fig. 3i; *Swsap1^-/-^ Chk2^-/-^, Chk2^-/-^*, n=1. Error bars, mean±s.d. ****, *P*≤0.0001; Mann-Whitney test, one-tailed. **(f)** *Swsap1^-/-^ Chk2^-/-^* spermatocytes that progress to mid-pachynema and beyond are increased in number relative to that observed in *Swsap1^-/-^* mice, as demonstrated by histone H1t staining. Total number of mid-pachytene, late-pachytene, and diplotene spermatocytes divided by the total number of spermatocytes analyzed from adult testes. Mice: Control, *Swsap1^-/-^*, from Fig. 2a; *Chk2^-/-^, Swsap1^-/-^ Chk2^-/-^*, n=2. *, *P*≤0.05; ****, *P*≤0.0001; Fisher’s exact test, two-tailed. **(g)** Spermatocytes in adult *Swsap1^-/-^ Chk2^-/-^* mice mostly arrest at pachynema, but tubules occasionally contain round and elongated spermatids (blue rectangles and insets). Sections were stained with H&E. Scale bars, 100 μm (top panel), 50 μm (bottom panel), and 20 μm (inset).

As Shu-mutant females are sterile, we also tested for evidence of HR defects in oocytes from embryonic day 18.5, when most have entered pachynema (Supplementary Fig. 4g). MLH1 foci are present in *Swsap1^-/-^* oocytes at mid-pachynema, but fewer as in spermatocytes (Fig. 3h,i). Furthermore, most mutant oocytes have ≥1 chromosome pair that is not synapsed and/or lacks an MLH1 focus (Supplementary Fig. 4h). Thus, the Shu complex is essential for crossover formation in both male and female meiosis.

Although some mutant oocytes form normal numbers of MLH1 foci, oocytes are mostly eliminated within a few days after birth (Supplementary Fig. 2e), resembling other DNA repair-defective mutants like *Dmc1^-/-^* (Ref.^22^). Oocyte loss in *Dmc1^-/-^* mice at 3 weeks can be partially rescued by eliminating the DNA damage checkpoint kinase CHK2, although these mice still lack primordial follicles and most oocytes are depleted in adult *Dmc1^-/-^ Chk2^-/-^* females (2-month-old)^32^. By contrast, *Swsap1^-/-^ Chk2^-/-^* adults show a complete rescue of ovary size (Fig. 4a,b and Supplementary Fig. 5a). Double mutant ovaries contain primordial follicles, unlike ovaries from *Swsap1^-/-^* mice, although the rescue is incomplete (Fig. 4c,d). Loss of CHK2, however, does not alleviate the synapsis defects (open circles; Fig. 4e) or improve MLH1 focus numbers (Fig. 4e and Supplementary Fig. 5b). Nonetheless, *Swsap1^-/-^ Chk2^-/-^* mutant females produce viable, fertile offspring, although with about half the litter size of controls (Supplementary Table 3b).

This rescue is remarkable, given the absence of MLH1 foci on one or more chromosomes (Supplementary Fig. 5b). CHK2 loss also suppresses infertility in females with a hypomorphic *Trip13* mutation^32^, although in this mutant MLH1 focus numbers in oocytes are inferred to be nearly normal (as reported for spermatocytes^33,34^). It would be interesting to determine whether viable pups from *Swsap1^-/-^ Chk2^-/-^* dams arise from oocytes with normal MLH1 focus counts or from fortuitous segregation of non-recombinant chromosomes^35^.

Unlike females, *Swsap1^-/-^ Chk2^-/-^* males exhibit only minimal rescue, with only marginally larger testes. Some tubules exhibit greater cellularity, H1t-positive spermatocytes are increased in number, and round and elongated spermatids are occasionally observed (Fig. 4a,f,g and Supplementary Fig. 5c). Nonetheless, tubules still mostly lack the full complement of germ cells (Fig. 4g) and mice remain infertile (Supplementary Table 3b). The minimal rescue by CHK2 loss in males could reflect a DSB-independent arrest tied to sex-body defects^29^.

Because some chromosomes in Shu-mutant meiocytes synapse and some cells progress to have MLH1 foci, we reasoned that another mediator protein(s) promotes some level of DSB repair in the absence of the Shu complex. One obvious candidate is BRCA2. Loss of BRCA2 abolishes assembly of both RAD51 and DMC1 into foci, such that spermatocytes do not progress past early pachynema^3^. Although expression of BRCA2 lacking the C-terminal domain in *Brca2^Δ27/Δ27^* mice^36^ causes an HR defect in somatic cells^37^, there is little discernible effect on RAD51 and DMC1 foci in early meiotic cells and cells progress to later prophase I stages (Fig. 5a-g). This truncated BRCA2 protein lacks C-terminal RAD51 and DMC1 interaction sites^6, 7, 9^, but contains several other RAD51 and DMC1 interaction sites which can promote their assembly into foci^8,10^ (Fig. 5d-g). *Brca2^Δ27/Δ27^* mice are fertile, although testes are smaller (Supplementary Fig. 5d), likely due to late meiotic prophase defects (C.M.A. and M.J., unpublished results).

**Figure 5:**
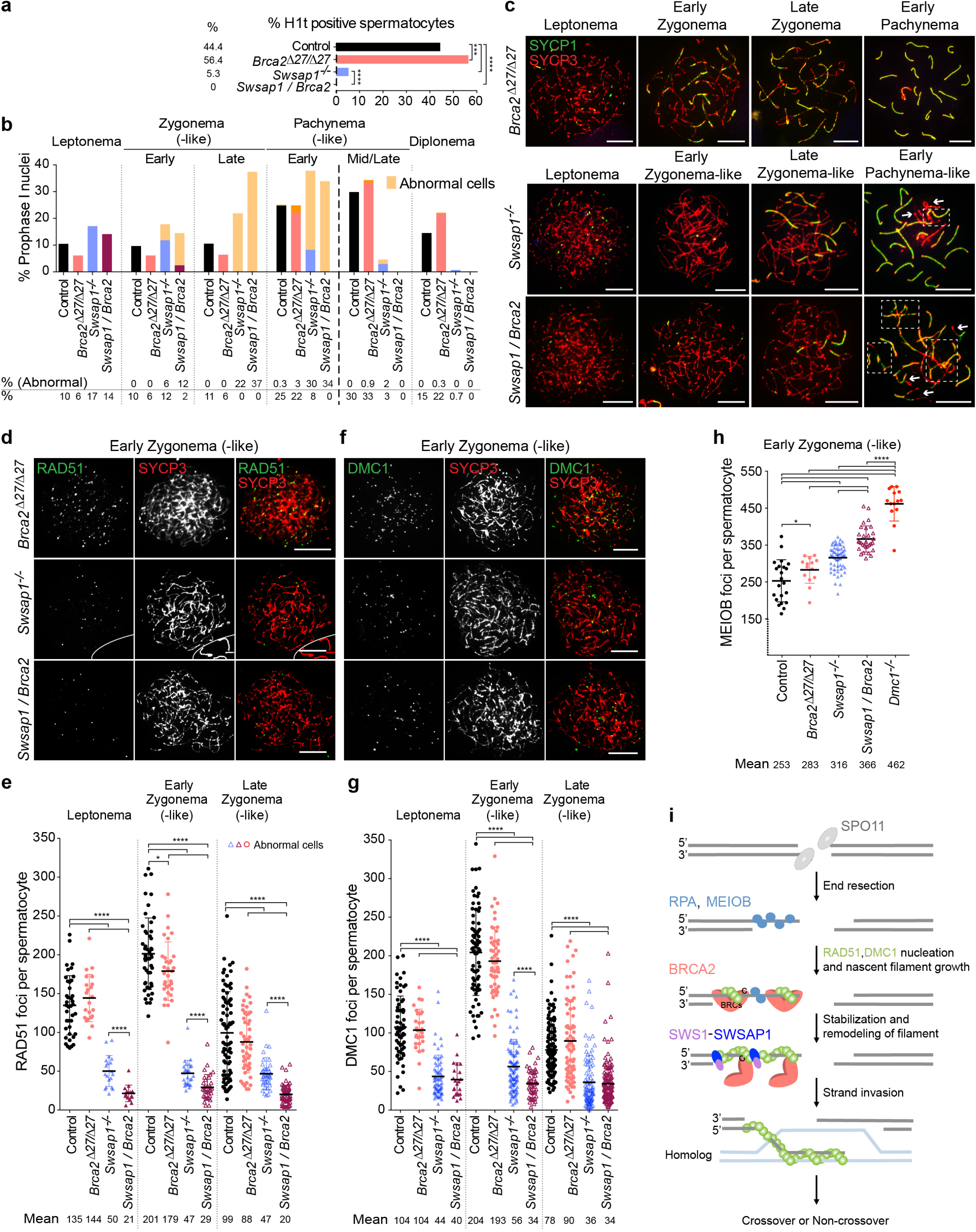
BRCA2 C terminus is dispensable for RAD51 and DMC1 focus formation early in meiosis but becomes critical when the Shu complex is absent. **(a)** *Swsap1^-/-^(+1) Brca2^Δ27/Δ27^* spermatocytes do not progress past early pachynema in contrast to *Swsap1^-/-^(+1)* spermatocytes in which a fraction progresses, as indicated by histone H1t staining. Total number of mid-pachytene, late-pachytene, and diplotene spermatocytes divided by the total number of spermatocytes analyzed from adult testes. Mice: Control, n=8 (combined with those from Fig. 2a); *Brca2^Δ27/Δ27^*, n=3; *Swsap1^-/-^*(+1) *Brca2^Δ27/Δ27^*, n=4; *Swsap1^-/-^(+1)*, n=4 (combined with those from Supplementary Fig. 3a). ***, *P*≤0.001; ****, *P*≤0.0001; Fisher’s exact test, twotailed. (b,c) *Swsap1^-/-^(+1) Brca2^Δ27/Δ27^* spermatocytes show altered meiotic progression and spermatocytes have aggravated synapsis defects relative to *Swsap1^-/-^* spermatocytes. Percentage of spermatocytes in the indicated stages in b with representative images in c. Most of the *Swsap1^-/-^(+1) Brca2^Δ27/Δ27^* early-zygotene (-like) cells have delayed synapsis, while this is only seen in a fraction of *Swsap1^-/-^(+1)* spermatocytes (light orange bars). By early pachynema, all double-mutant spermatocytes display incomplete homolog synapsis and/or nonhomologous synapsis. Boxes in early pachytene-like cells in c highlight non-homologous synapsis, and arrows indicate unsynapsed chromosomes. **(d-g)** RAD51 and DMC1 focus counts are further reduced in *Swsap1^-/-^(+1) Brca2^Δ27/Δ27^* spermatocytes relative to *Swsap1^-/-^(+1)*. Representative chromosome spreads at early zygonema (-like) are shown in **d,f** with focus counts for the indicated stages in **e,g**. Control, n=5 (RAD51, combined with 2 mice from Fig. 3e) and n=4 (DMC1, combined with 2 mice from Fig. 2g); *Swsap1^-/-^(+1)*, n=3 (RAD51, combined with that from Supplementary Fig. 3f) and n=4 (DMC1, combined with that from Supplementary Fig. 3g); *Brca2^Δ27/Δ27^, Swsap1^-/-^ (+1) Brca2^Δ27/Δ27^*, n=3 (RAD51) and n=4 (DMC1). **(h)** MEIOB focus counts at early zygonema (-like) are further increased in *Swsap1^-/-^ (+1) Brca2^Δ27/Δ27^* spermatocytes relative to *Swsap1^-/-^(+1)*, but still lower than that in *Dmc1^-/-^* spermatocytes. Mice: Control, n=3 (combined with those from Fig. 3b); *Swsap1^-/-^ (+1)*, n=3 (combined with that from Supplementary Fig. 4c); *Brca2^Δ27/Δ27^*, n=2; *Swsap1^-/-^(+1) Brca2^Δ27/Δ27^*, n=3; *Dmc1^-/-^*, n=1 (from Fig. 3b). **(i)** Model for the functional interaction between the mouse Shu complex and the BRCA2 C terminus during meiotic HR. While the Shu complex is critical for stable RAD51 and DMC1 nucleoprotein filament assembly during meiosis, the BRCA2 C terminus can provide some activity in the absence of this complex. Error bars in **e,g,h**: mean±s.d. Scale bars in **b,d,f**: 10μm. Each symbol in **e,g,h** is the total number of foci from a single nucleus. Solid symbols in **e,g,h**: normal cells. Open symbols in **e,g,h**: abnormal cells. *, *P*≤0.05 in **e,h**; **, *P*≤0.01 in **g**; ****, *P*≤0.0001 in **e,g,h**; Mann-Whitney test, one-tailed. See

We asked whether the BRCA2 C terminus plays a role in the absence of the Shu complex to support RAD51 and DMC1 foci and thus inter-homolog repair. Indeed, loss of the BRCA2 C terminus aggravates the homolog synapsis defects in *Swsap1^-/-^* mice, and H1t-positive cells are absent from *Swsap1^-/-^ Brca2^Δ27/Δ27^* mice, indicating a fully penetrant meiotic arrest (Fig. 5a-c). Testis weights are also slightly reduced (Supplementary Fig. 5d). Unlike *Swsap1^-/-^* spermatocytes, most double mutant cells at early zygonema show delayed synapsis, and all cells at early pachynema display asynapsis and/or nonhomologous synapsis, with fewer fully synapsed chromosomes (Fig. 5b,c and Supplementary Fig. 5e). Importantly, *Swsap1^-/-^ Brca2^Δ27/Δ27^* spermatocytes show even fewer foci at leptonema and early zygonema for both RAD51 (2.4 and 1.6-fold reduction, respectively) and DMC1 (1.1 and 1.6-fold reduction, respectively) compared with *Swsap1^-/-^* (Fig. 5d-g). There is a concomitant further increase in MEIOB foci (1.2-fold; Fig. 5h), although interestingly, MEIOB foci are still fewer than in *Dmc1^-/-^* spermatocytes. We conclude that, although the BRCA2 C terminus is largely dispensible for meiotic HR in the presence of SWSAP1, it functions in the absence of the Shu complex to support stable assembly of RAD51 and DMC1 on resected DNA ends. However, it is clearly not sufficient to overcome the loss of the Shu complex at most DSBs.

Strand invasion by RAD51 and DMC1 is a central step in meiotic HR. Our studies reveal a key requirement for the mouse Shu complex in stable assembly of sufficient numbers of both RAD51 and DMC1 nucleoprotein filaments to promote homolog pairing during meiosis. As a result, resected DNA ends accumulate in the *Sws1^-/-^* and *Swsap1^-/-^* mutants, although not to the level seen when DMC1 itself is absent. As in mouse, Rad51 focus formation is severely reduced in budding yeast Shu mutants during meiosis; however, our observations contrast with those from yeast, where Dmc1 focus formation is relatively unaffected^13^. Other yeast mediator proteins likely have Dmc1-specialized roles, in particular the Mei5-Sae3 complex^2^. Because of the embryonic lethality associated with mutation of canonical RAD51 paralogs^10^, it is unclear if or how these proteins contribute to mammalian meiosis. One hint comes from mice expressing a hypomorphic *Rad51c* allele, as prophase spermatocytes from these mice show reduced RAD51 focus formation, although DMC1 was not examined^38^. Thus, as in mitotic cells, multiple protein complexes are likely needed to promote recombinase activity in meiotic cells, although how they functionally interact remains to be elucidated.

We envision the mouse Shu complex stabilizing both RAD51 and DMC1 nucleoprotein filaments (Fig. 5i), and possibly remodeling them, as reported for Rad51 by Shu complexes and canonical RAD51 paralogs in other organisms^13, 39, 40^. By contrast, the primary role of BRCA2 is nucleation of RAD51 and DMC1 nucleoprotein filaments, a role ascribed to the BRC repeats in the center of the protein^8,10^. However, the BRCA2 C terminus has also been implicated in RAD51 filament stabilization by selectively binding to the interface between two RAD51 protomers^6,7^. It will be interesting to determine if the BRCA2 C terminus also promotes DMC1 filament stabilization. Thus, while the Shu complex and the BRCA2 C terminus have overlapping biochemical roles during meiotic HR, the Shu complex is clearly more critical given the infertility of Shu mutant mice.

## Online Methods

### Mouse care

The care and use of mice were performed in accordance with the Memorial Sloan Kettering Cancer Center (MSKCC) Institutional Animal Care and Use Committee guidelines.

### Generation and genotyping of Shu mutant mice

We targeted *Sws1* and *Swsap1* with TALE nucleases directed to each gene’s first exon, close to the translational start sites. TALEN pairs (RNA) were injected into fertilized mouse eggs, derived from superovulated CBA/J x C57BL/6J F1 females mated with C57BL/6J males, which were then implanted into pseudo-pregnant females^41,42^. To initially genotype founder mice, at least 10 cloned PCR products from each of 22 *Sws1* and 10 *Swsap1* founders were sequenced. Founders were backcrossed to C57BL/6J to separate multiple alleles and then further backcrossed for 3-6 additional generations prior to generating experimental mice.

Genotyping for *Sws1* was done by PCR-sequencing using the following PCR primers: Sws1-A: 5’-CCTGCAGGGCGCGTGAAGTTC, Sws1-B: 5’-ACCGGCTCGCACTCAGGGATC under the following conditions: 94 °C, 3 min; 35 cycles of 94 °C, 30 sec; 55 °C, 1 min and 65 °C, 30 sec; and a final extension of 72 °C, 5 min. The PCR product (259 bp) was sequenced using *Sws1*-A primer and sequencing reads were aligned against the wild-type controls to detect the 1-bp deletion.

Genotyping for *Swsap1* was done using the following PCR primers *Swsap1-C:* 5’-TCTGTGAACTATAGCCAATGAGGC, and Swsap1-D: 5’-AACTGTCACTCAGGCGCGAACTAG under the following PCR conditions: 94 °C, 3 min; 35 cycles of 94 °C, 30 sec; 55 °C, 1 min and 65 °C, 30 sec; and a final extension of 72 °C, 5 min. The *Swsap1(+1)* allele was genotyped by PCR-sequencing using the *Swsap1-C* primer; the *Swsap1Δ1A* allele was genotyped by running PCR products on a 2.4% agarose gel. The wild-type product is 396 bp and the mutant is 265 bp.

*Chk2* (Ref.^43,44^) and *Brca2^Δ27^* (Ref.^36,37^) mice and genotyping were previously described.

### RT-PCR

Twenty milligrams of mouse tissue was incubated with 1 ml Triazol and homogenized with a Dounce homogenizer. The extract was transferred to Eppendorf tubes and incubated for 5 min at room temperature. Extracts were centrifuged at 12,000 *g* for 10 min at 4 °C. Supernatants were transferred to another Eppendorf tube and RNA was extracted using chloroform followed by isopropanol precipitation. The RNA pellet was dissolved in H_2_O. To prepare the cDNA library, Superscript one-step RT-PCR kit was used (Invitrogen). To amplify cDNA for *Sws1* and *Swsap1*, the following primers were used: Sws1-RT-A: 5’-AAGTTCGCAGCGCCCGGG, *Sws1-* RT-B: 5’-CTAGGCTTCTGTCTTTGAAGTCC, Swsap1-RT-A: 5’-ATGGCGGAGGCGCTGAGG, Swsap1-RT-B: 5’-TCAGGTCTTTGAATCTGCACCTG. The following conditions were used for PCR: 94 °C, 2 min; 30 cycles of 94 °C, 1 min; 65 °C, 1 min and 72 °C, 1 min; and a final extension of 72 °C, 10 min. PCR products were separated on 1.0% agarose gels, excised, and DNA was purified and sequenced. The Sws1-RT-A and Swsap1-RT-A primers were used for sequencing.

### Histology

Ovaries and testes were dissected from animals at the stated ages and fixed in Bouin’s and stained with PAS, fixed in 4% PFA and stained with H&E, or fixed in 4% PFA and stained with hematoxylin and antibodies against DDX4/Vasa (Abcam, ab13840; 2.5 μg/ml) or c-Kit (Cell Signaling, 3074; 0.75 μg/ml) or were TUNEL-stained (Roche, 03333566001 and 11093070910). Staging of PAS- or H&E-stained testes sections was performed as described^18^. For follicle counts, ovaries were serially sectioned at 6 μm thickness, and follicles were counted in every fifth section, without further correction. The results were from one ovary from each animal.

### Spermatocyte chromosome spreads and immunofluorescence microscopy

Testes were collected from 2-4 month-old mice and spermatocytes were prepared for surface spreading and processed using established methods for immunofluorescence^45^, using the following primary antibodies in dilution buffer (0.2% BSA, 0.2% fish gelatin, 0.05% Triton X-100, 1xPBS), with incubation overnight at 4 °C: mouse anti-SYCP3 (Santa Cruz Biotechnology, sc-74569; 1:200), rabbit anti-SYCP3 (Abcam, ab15093; 1:500), goat anti-SYCP3 (Santa Cruz Biotechnology, sc-20845; 1:200), rabbit anti-SYCP1 (Novus, NB-300-229; 1:200), mouse anti-γH2AX (Millipore, 05-636; 1:500), rabbit anti-RAD51 (Calbiochem, PC130; 1:200), rabbit anti-DMC1 (Santa Cruz Biotechnology, sc-22768; 1:200), rabbit anti-MEIOB (kindly provided by P.J. Wang, University of Pennsylvania; 1:200), rat anti-RPA2 (Cell Signaling Technology, 2208S; 1:100), rabbit anti-MSH4 (Abcam, ab58666; 1:100), mouse anti-MLH1 (BD Biosciences, 51-1327GR; 1:50) and guinea pig anti-H1t (kindly provided by M.A. Handel, Jackson Laboratory; 1:500). Slides were subsequently incubated with the following secondary antibodies at 1:200 to 1:500 dilution for 1h at 37 °C: 488 donkey anti-mouse (Life Technologies, A21202), 488 donkey anti-rabbit (Life Technologies, A21206), 488 goat anti-rat (Life Technologies, A11006), A568 goat anti-mouse (Molecular probes, A-11019), 568 goat antirabbit (Life Technologies, A11011), 594 donkey anti-mouse (Invitrogen, A21203), 594 goat anti-rabbit (Invitrogen, A11012), donkey 594 anti-goat (Invitrogen, A11058), 647 donkey antimouse (Life Technologies, A31571), 647 donkey anti-rabbit (Invitrogen, A31573), 647 goat anti-guinea pig (Life Technologies, A21450). Cover slips were mounted with ProLong Gold antifade reagent with or without DAPI (Invitrogen, P36935 and P36934, respectively). Immunolabeled chromosome spread nuclei were imaged on a Marianas Workstation (Intelligent Imaging Innovations; Zeiss Axio Observer inverted epifluorescent microscope with a complementary metal-oxide semiconductor camera) using 100× oil-immersion objective. Images were processed using Image J for foci analysis and Photoshop (Adobe) to make the figures.

Spermatocytes were staged by assessing the extent of SYCP3 staining and synapsis (based on SYCP1 staining for Fig. 2, 5 and Supplementary Fig. 3). Only foci colocalizing with the chromosome axis were counted. In controls, leptotene cells are characterized by the presence of small stretches of SYCP3 and no SYCP1 staining. At early zygonema, homolog synapsis initiates, marked by the presence of SYCP1, and longer SYCP3 stretches are visible as chromosome axes continue to elongate. By late zygonema, chromosome axis formation completes at the same time that SYCP1 appears between homologs (>50% of overall synapsis). At pachynema, homologs are fully synapsed (co-localization of SYCP3 and SYCP1), except in the non-pseudoautosomal region of the XY chromosomes. Due to chromatin condensation at this stage, pachytene chromosome axes are shorter and thicker. H1t staining is used whenever possible to distinguish mid-/late-pachytene from early-pachytene cells. Late pachytene cells are further characterized by thickening of chromosome ends and elongation/curling of XY chromosomes. In diplotene cells, chromosome desynapsis ensues. *Sws1^-/-^* and *Swsap1^-/-^* cells at leptonema are indistinguishable from controls. Abnormal cells in mutants are characterized as follows: Early zygotene-like cells have chromosomes with long axes (SYCP3) and no SYCP1 stretches. Late zygotene-like cells have chromosomes with fully formed axes (SYCP3) and little or no synapsis in addition to elongated chromosomes completing synapsis as in controls. Early pachytene-like cells have unsynapsed chromosomes and/or incompletely synapsed homologs as well as non-homologous synapsis, but also display fully synapsed autosomes with thicker and shorter SYCP3 axes indicative of chromatin condensation characteristic of this stage. Midpachytene-like cells, which are H1t-positive, also display synaptic abnormalities and may contain chromosome fragments. Cells displaying normal homolog synapsis but chromosome end-to-end fusions are considered normal cells in control and mutants.

### Oocyte chromosome spreads

Prenatal ovaries were collected at embryonic day 18.5 and processed to obtain oocyte spreads as described^46^ with some modifications. Briefly, ovaries were placed in a 1.5 ml Eppendorf tube containing 0.7 ml isolation medium (TIM: 104 mM NaCl, 45 mM KCl, 1.2 mM MgSO_4_, 0.6 mM KH_2_PO_4_, 0.1% (w/v) glucose, 6 mM sodium lactate, 1 mM sodium pyruvate, pH 7.3, filter sterilized) and fragmented by pipetting up and down several times. After centrifuging for 3 min at 400 *g* and discarding the supernatant, 0.5 ml of 1 mg/ml collagenase (Sigma, C0130) in TIM was added to the ovarian fragments and tubes were incubated at 37 °C for 1 h with gentle shaking. Next, careful pipetting up and down was performed until large fragments of ovarian tissue were no longer visible. Following centrifugation for 3 min at 400 g, the supernatant was discarded, the pellet was carefully resuspended in 0.5 ml 0.05% trypsin (Sigma, T9935), and tubes were incubated at 37 °C for 5 min with gentle shaking. Then 0.5 ml DMEM containing 10% (v/v) FBS was added, carefully pippeted up and down, and centrifuged for 3 min at 400 g. The supernatant was discarded followed by the addition of 1 ml TIM to resuspend the cells by pipetting. Cells were again centrifuged for 3 min at 400 g, and after discarding the supernantant, cells were resuspended in 0.5 ml hypotonic solution (30 mM Tris-HCl pH 8.2, 50 mM sucrose, 17 mM Na-Citrate, 5 mM EDTA, 1× protease inhibitors), and incubated for 30 min to 1 h at room temperature. Positively charged, precleaned glass slides were placed in a humid chamber and a circle of ~1.0-1.5 cm diameter was marked in the center of the slide with a hydrophobic barrier pen; 40 μl of 1% (w/v) PFA containing 0.1% (v/v) Triton X-100 was placed into each circle and 10 μl cell suspension was slowly dropped on each slide within the PFA solution. Slides were slowly dried in a moist chamber that was closed for 2 h, then ajar for 30 min, and then open for 30 min. Slides were rinsed two times with milliQ H_2_O, and one time with 1:250 Photo-Flo 200 (Kodak, 1464510) solution. Slides were air-dried and stored at -80 °C. Staining of slides was then performed as for the spermatocyte chromosome spreads described above.

### Statistical analysis

Statistical analyses were performed using a Chi square test for animal breeding, a nonparametric two-tailed Mann-Whitney test for pup analysis, a two-tailed Student’s t-test for testis, ovary, and body weight comparisons, a two-tailed Fisher’s test for H1t cell analysis, a nonparametric one-tailed Mann-Whitney test for foci number comparisons, as a normal distribution could not be assumed, and an ANOVA test (Kruskal Wallis) for primordial follicle counts, due to the lack of follicles in the mutant. Error bars, mean±s.d.; ns, not significant; *, *P*<0.05; **, *P*<0.01; ***, *P*<0.001; ****, *P*<0.0001.

## Acknowledgements

We thank Mary Ann Handel and P. Jeremy Wang for antibodies, Kara Bernstein (University of Pittsburgh) and members of the Jasin and Keeney labs for discussions and critical reading of the manuscript, and Katia Manova and members of the MSKCC Molecular Cytology core facility for technical help. This work was supported by MSK Cancer Center Support Grant/Core Grant (NIH P30CA008748), NIH F32GM110978 (R.P.), BFU2016-80370-P (I.R.), R35 GM118092 (S.K.), R35 GM118175 (M.J.), and R01CA185660 (M.J.).

## Author’s contributions

C.M.A., R.P., P.J.R., S.K., and M.J. designed experiments. C.M.A., R.P., and P.J.R. performed experiments. I.R., S.K., and M.J. supervised the research. C.M.A., R.P., S.K., and M.J. wrote the paper with input from I.R.

## Competing financial interests

The authors declare no competing financial interests.

## Supplementary Figure legends

**Supplementary Figure 1:**
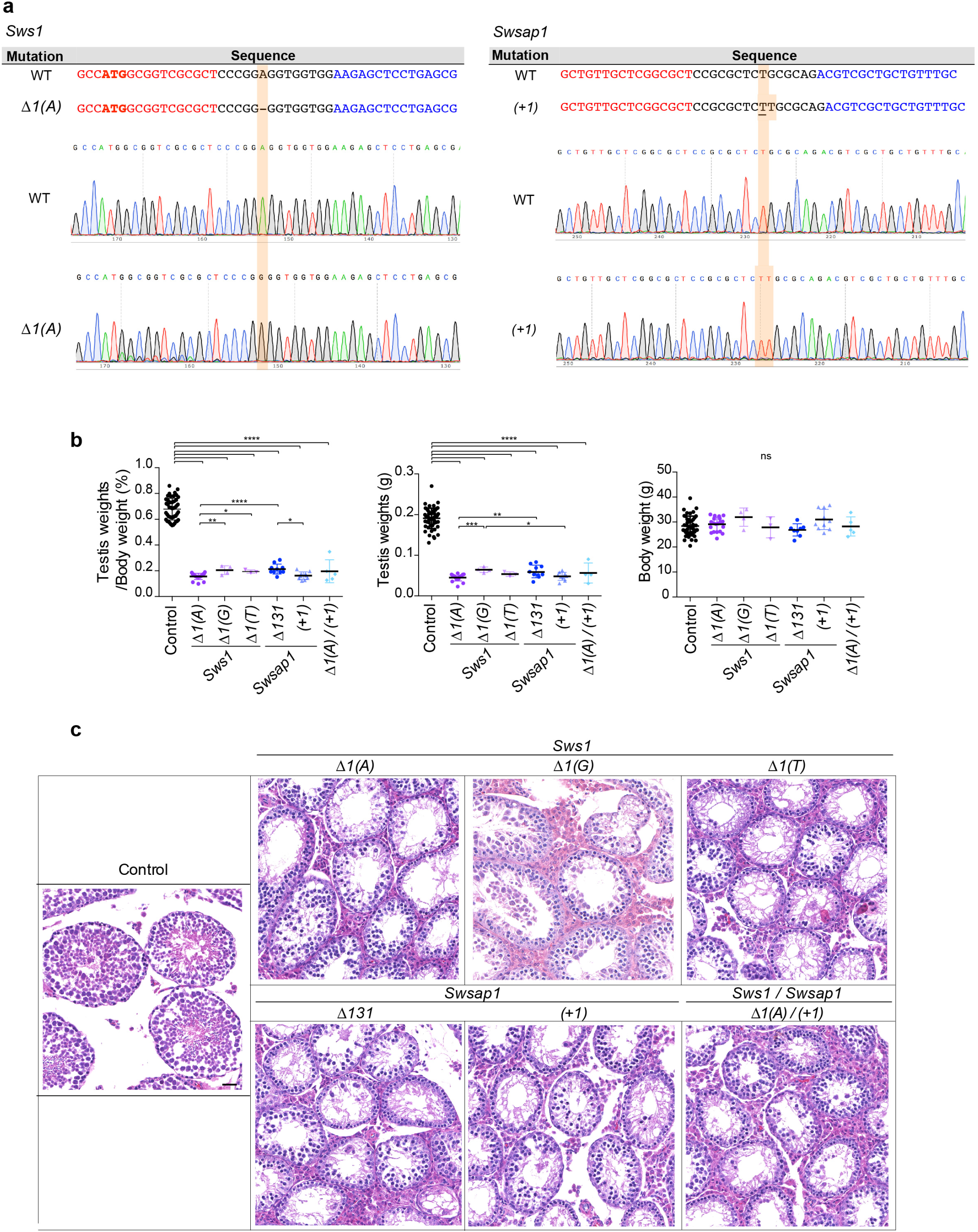
Meiotic defects in *Sws1* and *Swsap1* mutant testes. **(a)** RT-PCR was performed to confirm mutant alleles, as specific antibodies for SWS1 and Swsap1 could not be confirmed by western blot analysis. Sanger sequencing of cDNA from testis of *Sws1*Δ*1(A)* and *Swsap1(+1)* mice demonstrates the expected 1-bp deletion and 1-bp addition, respectively, in transcripts from these alleles. Positions of these mutations are highlighted in orange. n=2. **(b)** All Shu complex single and double mutants have similarly reduced testis to body weight ratios and testis weights but not body weights. The testis to body weight ratios are from the combined graph in Fig. 1c but here are separated according to allele. Mice: Control, n=45; *Sws1^-/-^* [Δ*1(A)*, n=20; Δ*1(G)*, n=4; Δ*1(T)*, n=3]; *Swsap1^-/-^* [Α*131*, n=10; *(+1)*, n=8]; *Sws1^-/-^ Swsap1^-/-^* (Δ*1(A)* / *(+1))*, n=5. *, *P*≤0.05; **, *P*≤0.01; ***, *P*≤0.001; ****, *P*≤0.0001; Student’s t-test, two-tailed. **(c)** Spermatocytes from single and double mutants show meiotic arrest. Testis sections were stained with H&E. Scale bars, 100 μm. n≥2.

**Supplementary Figure 2:**
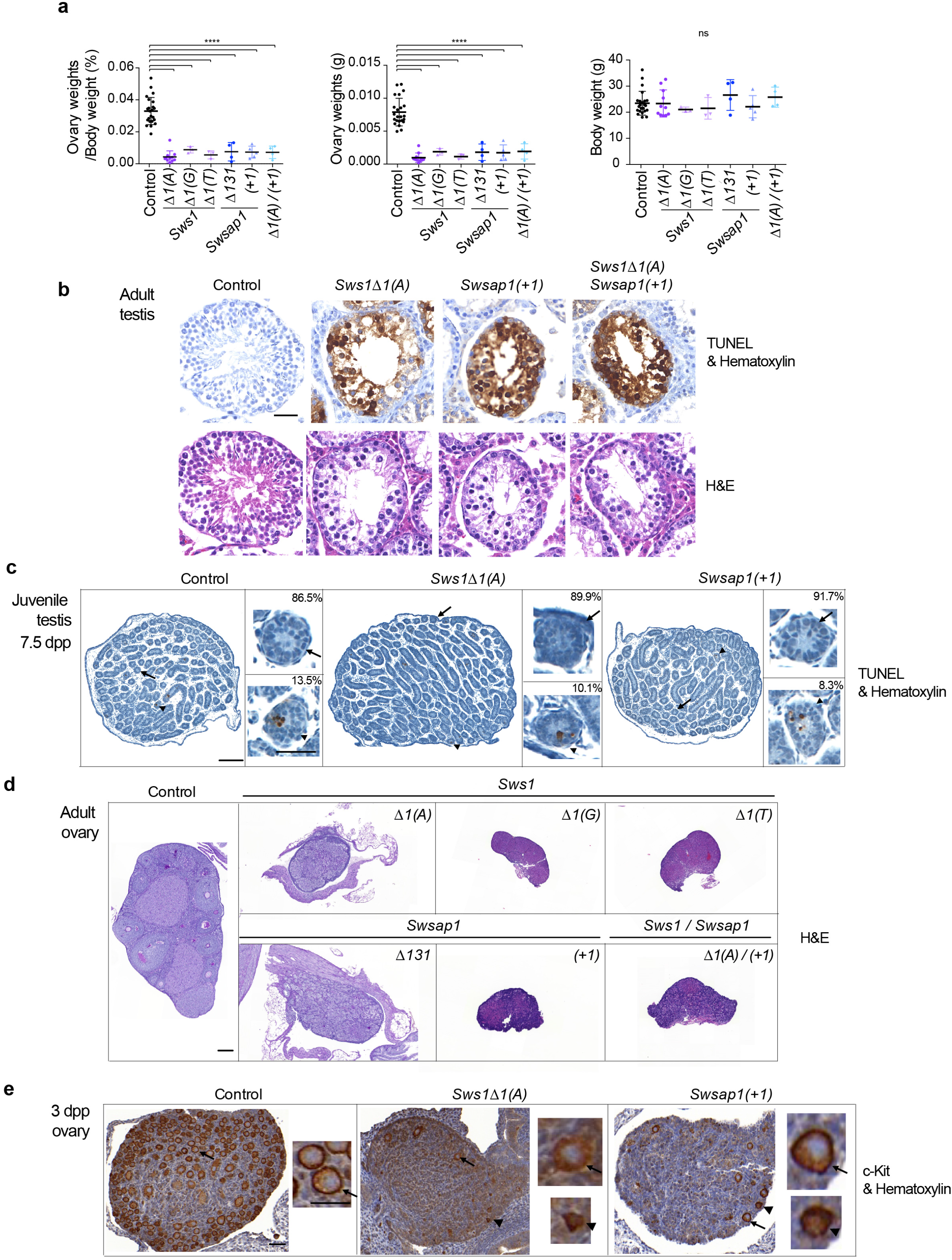
Shu mutants have abnormal ovaries and adult testes. **(a)** Shu complex single and double mutants have reduced ovary to body weight ratios and ovary weights but not body weights. The ovary to body weight ratios are from the combined graph in Fig. 1d but here are separated according to allele. Mice: Control, n=23; *Sws1^-/-^* [Δ*1(A)*, n=11; Δ*1(G)*, n=3; Δ*1(T)*, n=3]; *Swsap1^-/-^* [Δ131, n=4; *(+1)*, n=5]; *Sws1^-/-^ Swsap1^-/-^* (Δ*1(A)* / *(+1))*, n=4. ns, not significant compared to control; *P*≤0.0001, significant compared to control; Student’s t-test, two-tailed. **(b)** Widespread apoptosis is observed in seminiferous tubules from single- and double-mutant testes in adults. Sections were stained with hematoxylin and TUNEL was used to visualize apoptotic spermatocytes. Scale bar, 50 μm. n=2. **(c)** Apoptosis in 7.5-dpp *Sws1^-/-^* and *Swsap1^-/-^* juvenile testes is rarely observed indicating that premeiotic stages are not grossly affected. Arrowheads point to rare apoptotic cells in both control and Shu mutants. Insets indicate the % TUNEL-negative (arrows) and positive (arrowheads) tubules. Sections were stained and analyzed as in b. Scale bar, 50 μm and 20 μm insets. n=2. **(d)** Adult ovaries from single and double mutants show an absence of follicles at all stages of oocyte development. Sections were stained with H&E. Scale bar, 500 μm. n≥2. **(e)** Ovaries from 3-dpp *Sws1^−^* and *Swsap1^−^* mice show significantly reduced primordial and primary follicles, some of which appeared to be apoptotic based on the condensed chromatin. Insets show non-apoptotic (arrows) and possibly apoptotic (arrowheads) follicles. Sections were stained with c-Kit. Scale bar, 100 μm and 20 μm insets. n=2.

**Supplementary Figure 3:**
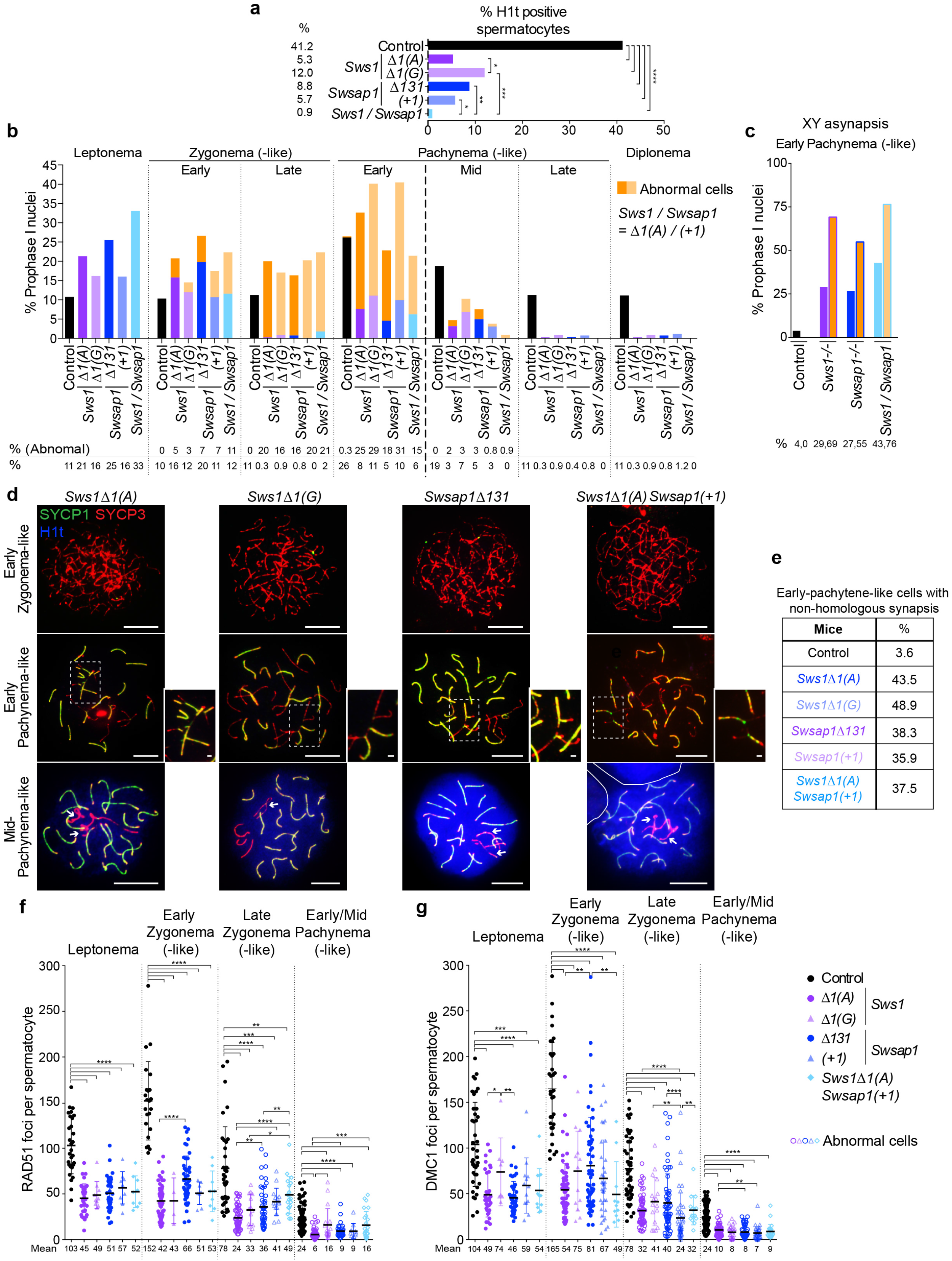
Spermatocytes from Shu complex mutant mice have defects in meiotic progression. **(a)** Histone H1t staining indicates that few spermatocytes in *Sws1* and *Swsap1* single and double mutants progress to mid-pachynema and beyond. Total number of mid-pachytene, late-pachytene, and diplotene spermatocytes divided by the total number of spermatocytes analyzed from adult testes. Mice: Control, *Sws1^-/-^*Δ*1(A), Swsap1^-/-^*Δ*131*, from Fig. 2a; *Sws1^-/-^*Δ*1(G), Sws1^-/-^ Swsap1^-/-^* (Δ*1(A) / (+1))*, n=1; *Swsap1^-/-^ (+1)*, n=2. *, *P*≤0.05; **, *P*≤0.01; ***, *P*≤0.001; ****, *P*≤0.0001; Fisher’s exact test, two-tailed. **(b-e)** *Sws1* and *Swsap1* mutants show altered meiotic progression and abnormal synapsis of both autosomes and sex chromosomes. Percentage of spermatocytes in each of the indicated meiosis prophase I stages is shown in **b**, with the percentage of early pachytene (-like) spermatocytes with XY asynapsis in **c**. Mice are the same as those in a, but combined for each mutant. Spermatocytes were staged as in Fig. 2b. The subset of cells with autosomal synapsis defects are summarized with the orange bars. Representative chromosome spreads of abnormal spermatocytes at early zygonema, and early and mid-pachynema are shown in **d**; percentages of early-pachytene cells with non-homologous synapsis are in **e**. Boxes and respective insets at early pachynema highlight non-homologous synapsis that is often in combination with homologous synapsis and partial asynapsis, resulting in chromosome tangles; arrows at mid-pachynema indicate unsynapsed chromosomes. Spermatocytes with complete homolog synapsis but chromosome end-to-end fusions are considered normal for this analysis. Scale bars in zoomed-out images, 10 μm. Scale bars in insets, 1 μm. **(f,g)** RAD51 and DMC1 focus counts are reduced in *Sws1* and *Swsap1* mutant spermatocytes. Each symbol is the total number of foci from a single nucleus. Solid symbol, normal cells. Open symbol, abnormal cells. Error bars, mean±s.d. *, *P*≤0.05; **, *P*≤0.01; ***, *P*≤0.001; ****, *P*≤0.0001; Mann-Whitney test, one-tailed. Mice: Control, *Sws1^-/-^*Δ*1(A), Swsap1^-/-^*Δ*131*, from Fig. 2e,g; *Sws1^-/-^*Δ*1(G), Swsap1^-/-^ (+1), Sws1^-/-^ Swsap1^-/-^* (Δ*1(A) / (+1))*, n=1.

**Supplementary Figure 4:**
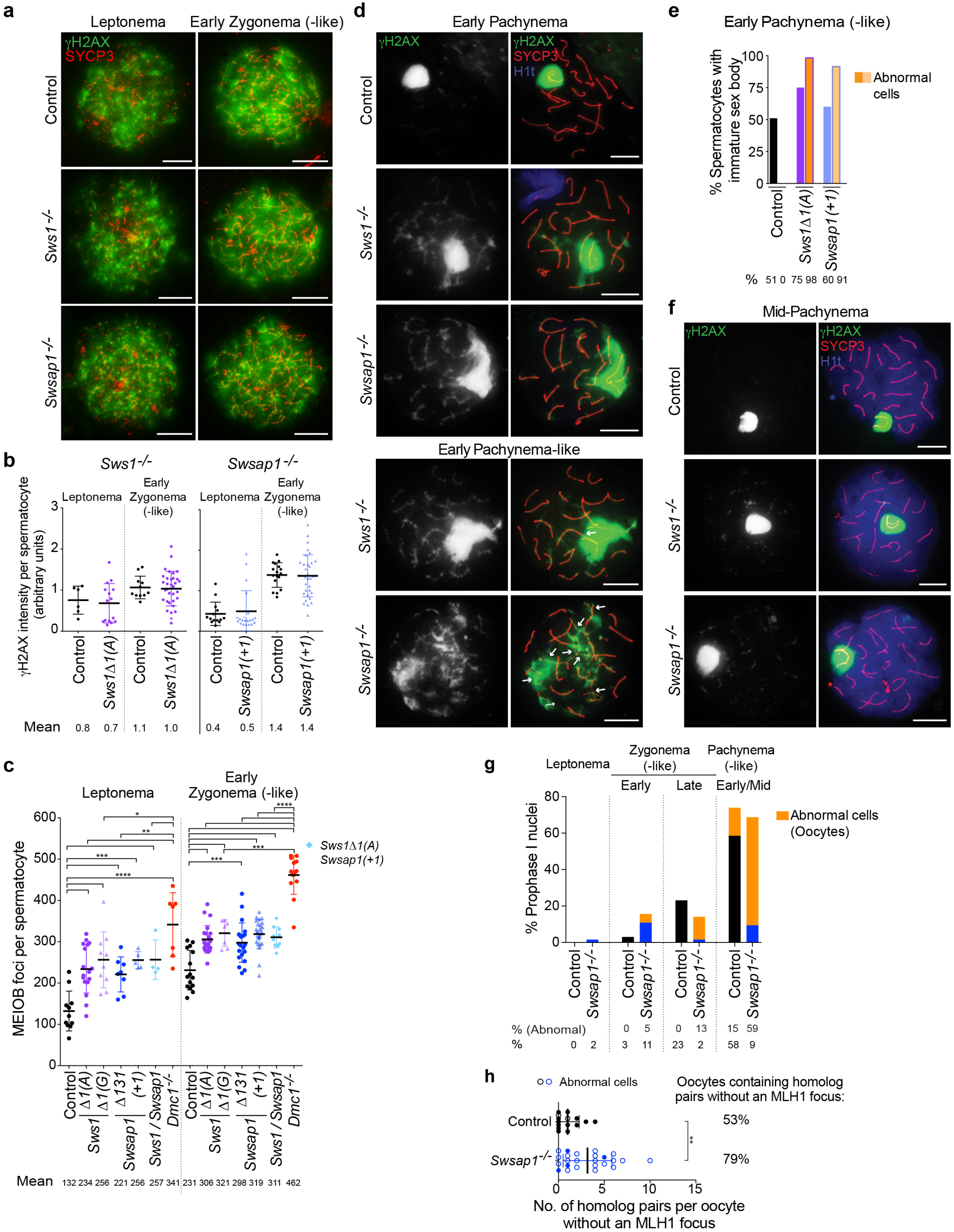
DSB formation appears to be normal in *Sws1* and *Swsap1* mutant spermatocytes, while DSB repair is compromised. **(a,b)** DSB formation, as measure by γH2AX levels, is unaffected in *Sws1^-/-^* and *Swsap1^-/-^* spermatocytes. Representative chromosome spreads at the indicated stages from adult mice in **a** with γH2AX integrated density in **b**. Scale bars, 10 μm. Each symbol in b is from a single nucleus to which the average background signal from four regions around each nucleus was subtracted. Error bars, mean±s.d. Mice: Control, *Sws1^-/-^*Δ*1(A), Swsap1^-/-^(+1)*, n=1. **(c)** MEIOB focus counts are increased in *Sws1^-/-^* and *Swsap1^-/-^* spermatocytes, but not as high as in *Dmc1^-/-^* spermatocytes. Each symbol is the total focus number from a single nucleus. Solid symbol, normal cells. Open symbol, abnormal cells. Error bars, mean±s.d. Mice: Control, *Sws1^-/-^* Δ*1(A), Swsap1^-/-^*Δ*131, Dmc1^-/-^*, from Fig. 3b; *Sws1^-/-^*Δ*1(G), Swsap1^-/-^(+1), Sws1^-/-^ Swsap1^-/-^* (Δ*1(A) / (+1))*, n=1. *, *P*≤0.05; **, *P*≤0.01; ***, *P*≤0.001; ****, *P*≤0.0001; Mann-Whitney test, one-tailed. **(d-f)** At early pachynema, *Sws1^-/-^* and *Swsap1^-/-^* spermatocytes show unrepaired DSBs associated with defects in sex body formation/maturation, whereas the small fraction of mid-pachytene mutant spermatocytes are apparently more repair proficient. Representative chromosome spreads at the indicated stages in **d,f** with percentage of nuclei displaying immature sex body in **e**. Arrows indicate unsynapsed chromosomes. Scale bars, 10 μm. Control, n=3; *Sws1^-/-^* Δ*1(A), Swsap1^-/-^(+1)*, n=2. **(g,h)** Most oocytes from control and *Swsap1* mice at embryonic day 18.5 have entered pachynema with the majority of mutant oocytes showing synapsis defects and one or more bivalents without an MLH1 focus. Percentage of oocytes at each of the indicated stages in **g** with the number of homolog pairs that are not synapsed and/or are lacking an MLH1 focus in **h**. The total percentage of oocytes containing homolog pairs without an MLH1 focus is also indicated. Error bars in **h**, mean±s.d. Solid circles, oocytes with fully synapsed homologs. Open circles, abnormal oocytes containing homologs partially synapsed and/or one or two homolog pairs fully unsynapsed. Mice from Fig. 3i. **, *P*≤0.01; Mann-Whitney test, one-tailed.

**Supplementary Figure 5:**
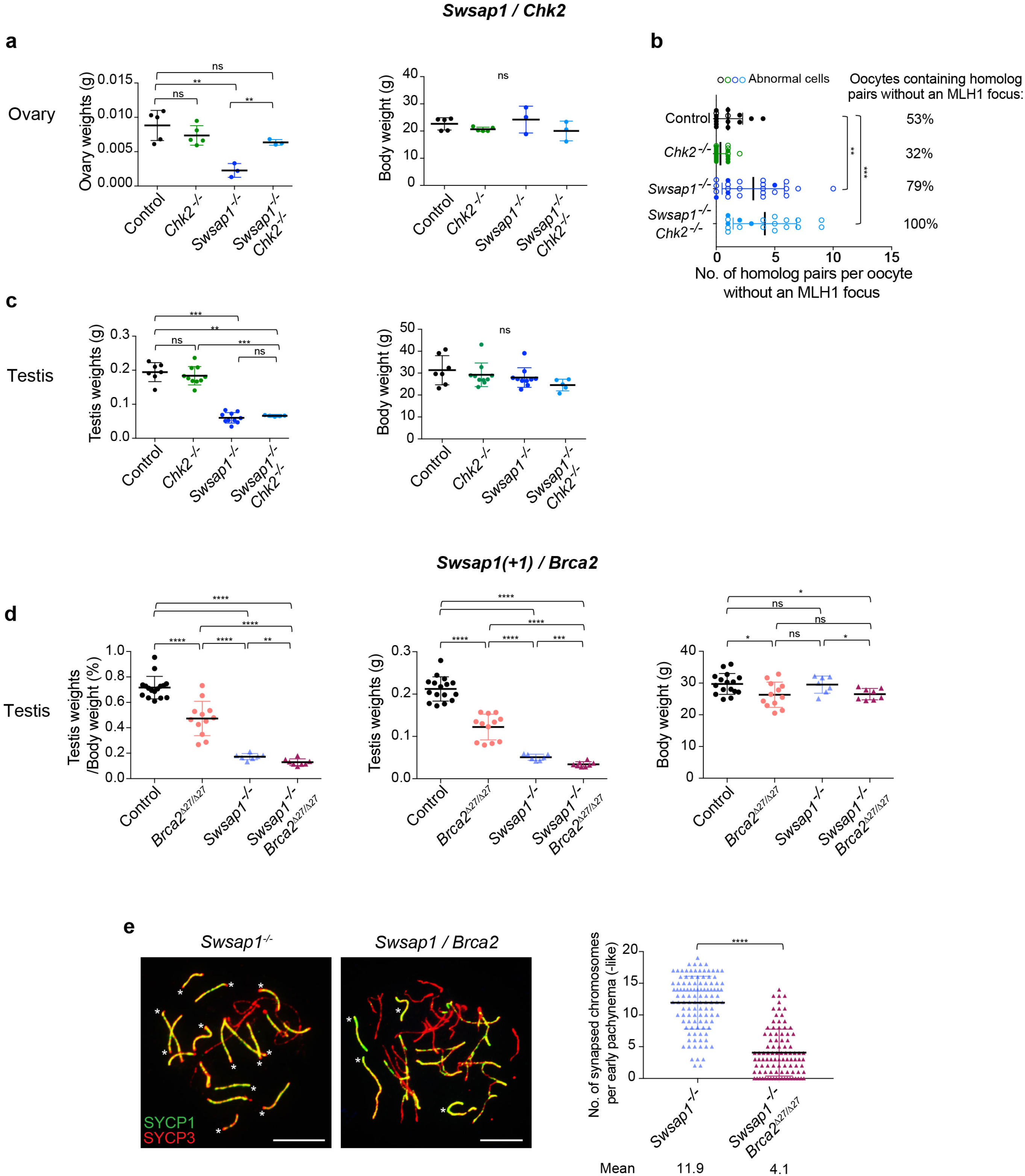
Effect of *Chk2* and *Brca2*^Δ*27*^ mutations on ovary and/or testis weights. **(a)** Ovary weights of *Swsap1^-/-^ Chk2^-/-^* mice are significantly higher than those of *Swsap1^-/-^* mice with no significant difference in the body weight. Mice: Control, n=5; *Chk2^-/-^*, n=5; *Swsap1^-/-^*, n=3; *Swsap1^-/-^ Chk2^-/-^*, n=3. ns, not significant; **, *P*≤0.01; Student’s t-test, two-tailed. **(b)** All *Swsap1^-/-^* and *Swsap1^-/-^ Chk2^-/-^* oocytes from mice at embryonic day 18.5 have one or more homologs fully unsynapsed and/or lacking one MLH1 focus. The total percentage of oocytes containing homologs without an MLH1 focus is also indicated. Solid circles, oocytes with homologs fully synapsed. Open circles, abnormal oocytes with homologs partially synapsed and/or one or more homolog pairs fully unsynapsed. Mice from Fig. 4e. Data for control and *Swsap1^-/-^* are from Supplementary Fig. 4h. **, *P*≤0.01; ***, *P*≤0.001; Mann-Whitney test, one-tailed. **(c)** Testis weights and body weights of *Swsap1^-/-^ Chk2^-/-^* and *Swsap1^-/-^* mice are similar. Mice: Control, n=7; *Chk2^-/-^*, n=10; *Swsap1^-/-^*, n=10; *Swsap1^-/-^ Chk2^-/-^*, n=5. **, *P*≤0.01; ***, *P*≤0.001; Student’s t-test, two-tailed. **(d)** Testis to body weight ratios and testis weights of *Swsap1^-/-^(+1) Brca2*^Δ*27*/Δ*27*^ are 1.3- and 1.7-fold reduced respectively compared to *Swsap1^-/-^(+1)* mice but no major difference in the body weights is seen in these mice. Mice: Control, n=16; *Brca2*^Δ*27*/Δ*27*^, n=12; *Swsap1^-/-^* (+1), n=7; *Swsap1^-/-^*(+1) *Brca2*^Δ*27*/Δ*27*^, n=8. *, *P*≤0.05; **, *P*≤0.01; ***, *P*≤0.001; ****, *P*≤0.0001; Student’s t-test, two-tailed. **(e)** The number of synapsed chromosomes in *Swsap1^-/-^(+1) Brca2*^Δ*27*/Δ*27*^ mutants is 3-fold reduced compared to *Swsap1^-/-^(+1)* spermatocytes. Representative chromosome spreads from early pachytene-like cells are shown. Scale bars, 10 μm. n=3. ****, *P*≤0.0001; Mann-Whitney, one-tailed. Error bars in **a-e**: mean±s.d.

---

**Supplementary Table 1 legend:**
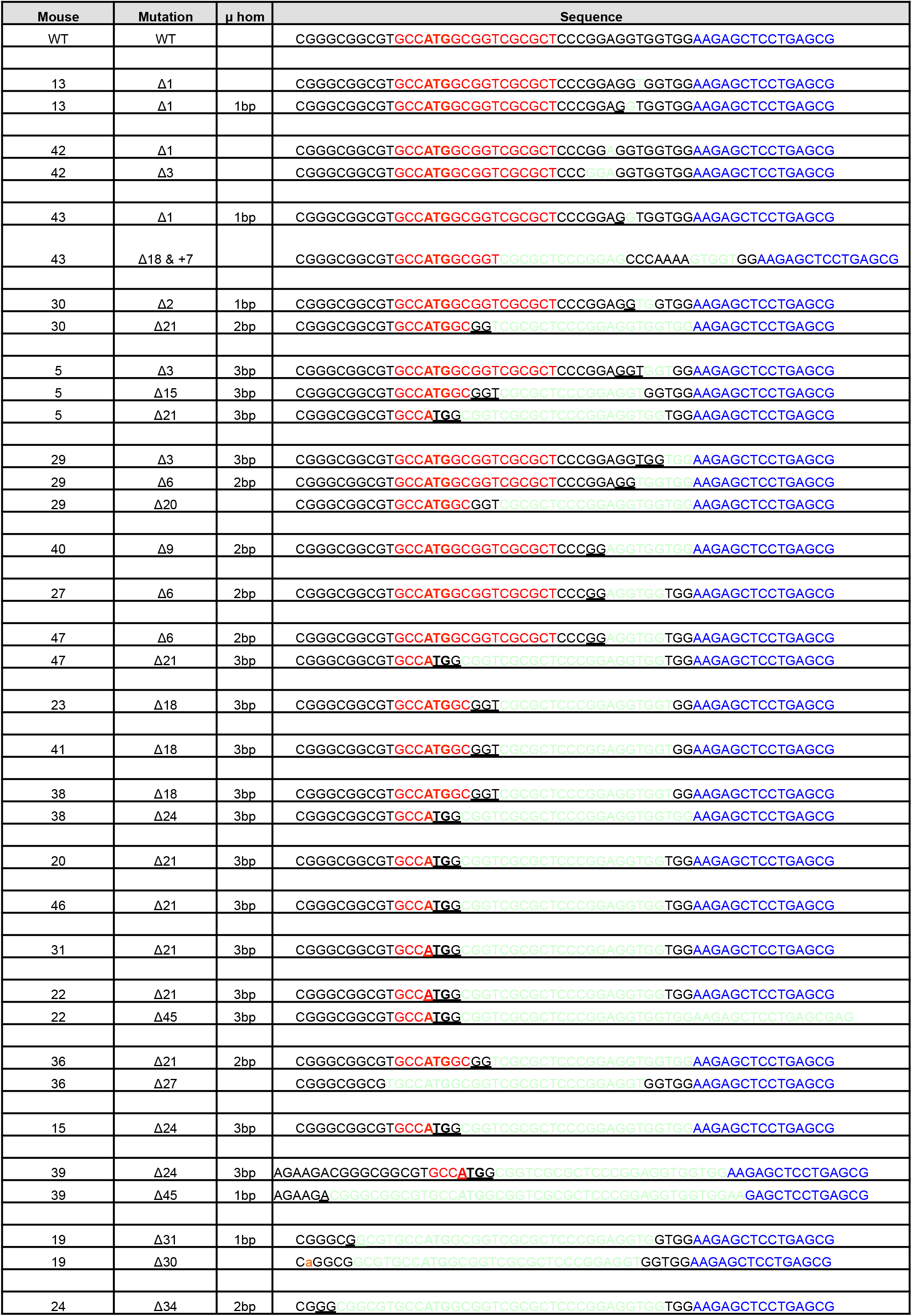
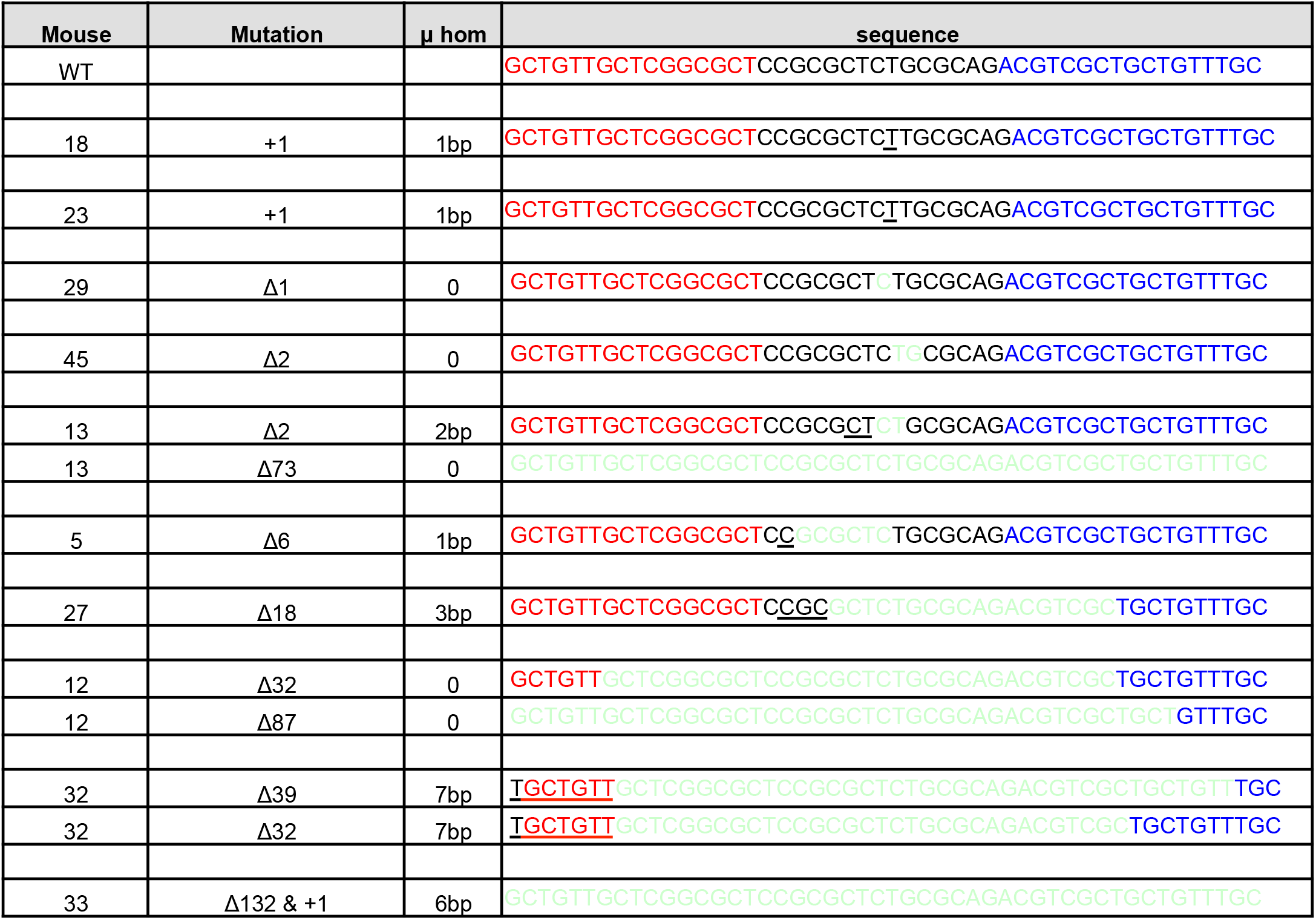
*Sws1* and *Swsap1* founder mice obtained after TALEN expression in **a** and **b,** respectively. Left and right TALEN binding sites are shown in red and blue. Indels are highlighted in light green, μhom refers to the microhomology at the breakpoint junction (underlined). Each *Sws1* founder mouse contained a wild-type allele except for 15, 20, 23, 27, 43 and 46. Each *Swsap1* founder mouse contained a wild-type allele. Forty-seven potential founders were born for *Sws1*, of which 22 were further analyzed based on T7 assays; 20 different indels were identified in these founder mice, of which 6 were frame-shift mutations. Forty-eight potential founders were born for *Swsap1*, of which 10 were further analyzed; 12 different indels were identified, of which 8 were frame-shift mutations.

**Supplementary Table 2 legend:** *Sws1^-/-^, Swsap1^-/-^* and double mutant mice are viable. Heterozygous mice for each genotype were bred to obtain homozygous knockouts. P-values were obtained using Chi squared analysis. Although *Sws1*Δ*1(A) and Swsap1(+1)* mutants appeared to be underrepresented in **a,** they were represented at the normal Mendelian ratio in **b.**

|  | Mutant allele | No. of mice | +/+ | +/- | -/- | P-values |
| --- | --- | --- | --- | --- | --- | --- |
| Sws1 | Sws1 Δ1(A) | 256 | 86 (34%) | 118 (46%) | 52 (20%) | 0.0050 |
|  | Sws1 Δ1(G) | 174 | 48 (28%) | 84 (48%) | 42 (24%) | 0.7332 |
|  | Sws1 Δ1(T) | 51 | 11 (22%) | 29 (56%) | 11 (22%) | 0.6185 |
| Swsap1 | Swsap1 Δ131 | 215 | 53 (24%) | 107 (50%) | 55 (26%) | 0.9793 |
|  | Swsap1 (+1) | 262 | 77 (29%) | 138 (53%) | 47 (18%) | 0.0222 |

b
|  | Control |  |  |  | Sws1<sup>-/-</sup> |  | Swsap1<sup>-/-</sup> |  | Sws1<sup>-/-</sup><br>Swsap1<sup>-/-</sup> |
| --- | --- | --- | --- | --- | --- | --- | --- | --- | --- |
| Sws1 / Swsap1 | +/+;<br>+/+ | Δ1(A)/+;<br>+/+ | +/+;<br>(+1)/+ | Δ1(A)/+;<br>(+1)/+ | Δ1(A)/Δ1(A);<br>+/+ | Δ1(A)/Δ1(A);<br>(+1)/+ | +/+;<br>(+1)/(+1) | Δ1/+;<br>(+1)/(+1) | Δ1(A)/Δ1(A);<br>(+1)/(+1) |
| Observed | 20<br>(7.1%) | 37<br>(13.1%) | 34<br>(12.0%) | 78<br>(27.7%) | 16<br>(5.7%) | 36<br>(12.8%) | 15<br>(5.3%) | 33<br>(11.7%) | 13<br>(4.6%) |
| Expected | 6.25% | 12.50% | 12.50% | 25.00% | 6.25% | 12.5% | 6.25% | 12.50% | 6.25% |

n=282

**Supplementary Table 3:**
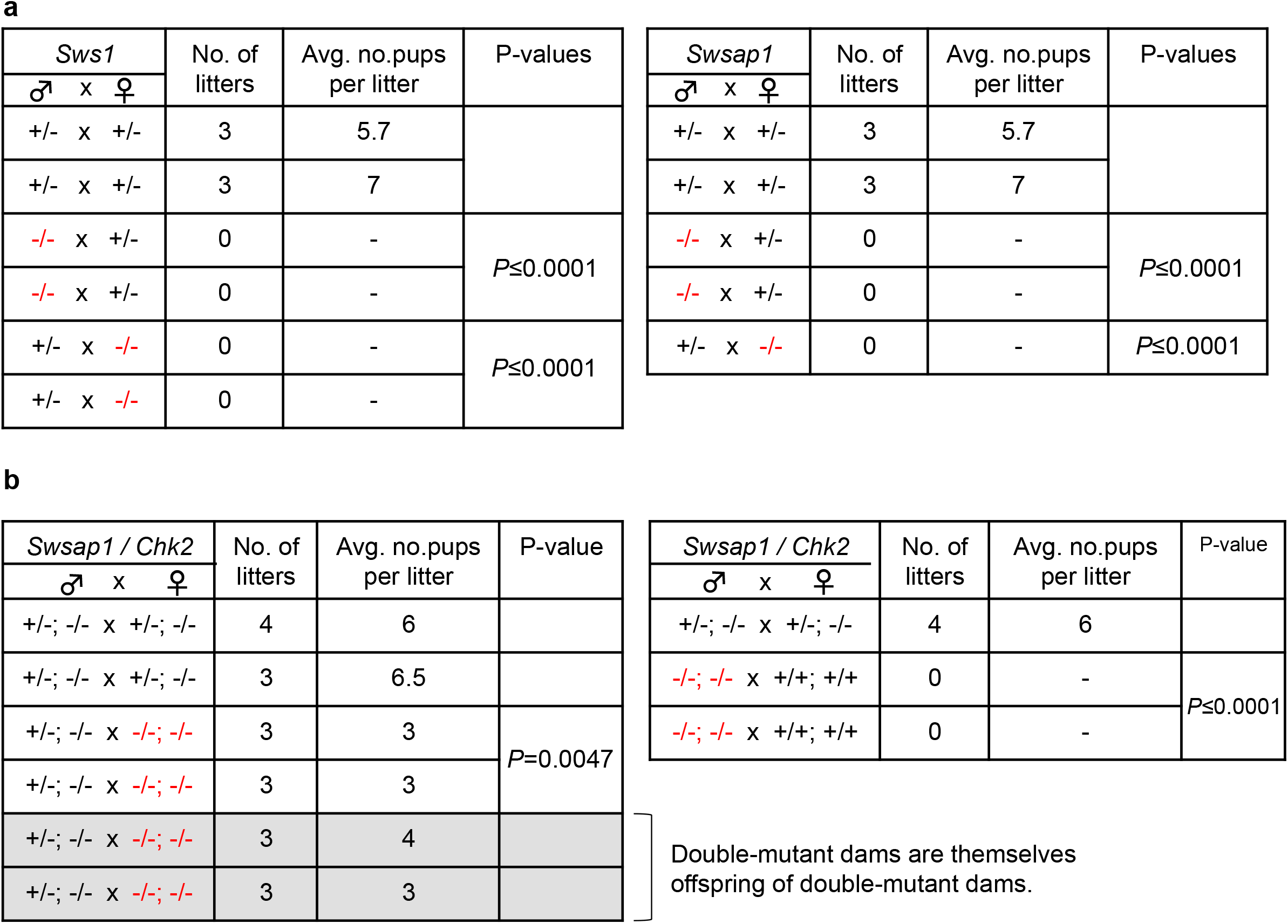
Fertility assessment of *Sws1^-/-^* and *Swsapt^A^* in **a** and *Swsap1^-/-^ Chk2^-/-^* in **b**. The single- or double-mutant mice (shown in red) were bred with heterozygous or wild-type mice for 4-5 months. Control heterozygous breedings were set up for the same period of time. P-values were obtained using Mann-Whitney test, two-tailed.

